# Focal adhesion-generated cues in extracellular matrix regulate cell migration by local induction of clathrin-coated plaques

**DOI:** 10.1101/493114

**Authors:** Delia Bucher, Markus Mukenhirn, Kem A. Sochacki, Veronika Saharuka, Christian Huck, Chiara Zambarda, Justin W. Taraska, Elisabetta Ada Cavalcanti-Adam, Steeve Boulant

## Abstract

Clathrin is a unique scaffold protein, which forms polyhedral lattices with flat and curved morphology. The function of curved clathrin-coated pits in forming endocytic structures is well studied. On the contrary, the role of large flat clathrin arrays, called clathrin-coated plaques, remains ambiguous. Previous studies suggested an involvement of plaques in cell adhesion. However, the molecular origin leading to their formation and their precise functions remain to be determined. Here, we study the origin and function of clathrin-coated plaques during cell migration. We revealed that plaque formation is intimately linked to extracellular matrix (ECM) modification by focal adhesions (FAs). We show that in migrating cells, FAs digest the ECM creating extracellular topographical cues that dictate the future location of clathrin-coated plaques. We identify Eps15 and Eps15R as key regulators for the formation of clathrin-coated plaques at locally remodelled ECM sites. Using a genetic silencing approach to abrogate plaque formation and 3D-micropatterns to spatially control the location of clathrin-coated plaques, we could directly correlate cell migration directionality with the formation of clathrin-coated plaques and their ability to recognize extracellular topographical cues. We here define the molecular mechanism regulating the functional interplay between FAs and plaques and propose that clathrin-coated plaques act as regulators of cell migration promoting contact guidance-mediated collective migration in a cell-to-cell contact independent manner.

## Introduction

Each cell is surrounded by a plasma membrane composed of lipids that separates the intracellular milieu from the extracellular space. To sense and interact with the extracellular environment, cells exploit both integral and peripheral membrane proteins and establish defined domains at the plasma membrane to mediate processes like signalling, endocytosis and mechanotransduction.

The main component of the environment surrounding cells is the extracellular matrix (ECM), a complex network of secreted macromolecules such as collagen, laminin and fibronectin^1,2^. Specific binding sites on these proteins serve as ligands for cellular receptors^3–5^. The most important of adhesion receptors are integrins, which, together with adaptors, scaffold and signalling proteins, assemble into the multi-layered adhesive unit referred to as focal adhesions (FAs)^3,6,7^. FAs mediate adhesion and signalling between the inside and the outside of the cell^7,8^. They are also viewed as mechanotransducing units coupling biophysical properties of the ECM to intracellular processes^8^. Indeed, not only the chemical but also the topographical and mechanical properties of the ECM influence cellular functions such as guiding cell migration^9,10^ and maintaining stem cell niches^11,12^. The ECM is not a static scaffold; it is a highly dynamic mesh constantly renewed and remodelled by cells, which in return react to the new properties of their surrounding matrix^13,14^. ECM remodelling is achieved both though cellular-mediated physical forces^15^ as well as through enzymatic digestions by specific enzymes like the matrix metalloproteinases (MMPs)^13^.

Beside FA, other plasma membrane-associated supramolecular complexes have been described to have mechanotransduction properties e.g. podosomes^16^ and cadherin-mediated cell-cell junctions^17^. Recently, plasma membrane-associated clathrin structures have been shown to assemble at specific plasma membrane sites in response to unique topographical profiles and mechanical properties of the extracellular environment^18–21^. Together with numerous adaptors and accessory proteins, clathrin molecules assemble at the plasma membrane to form a highly dynamic array^22,23^. During clathrin-mediated endocytosis (CME), small transient clathrin coats can form initially as curved or flat arrays which will rearrange to form clathrin-coated pits (CCPs) with a typical diameter of 100-150 nm^24–26^. Distinct from these small endocytic structures, flat long-lived larger clathrin coats, known as clathrin-coated plaques, are frequently observed^27^. These clathrin-coated plaques appear to have pleiotropic functions. They can facilitate endocytosis and signalling *via* clustering of plasma membrane receptors and nucleating endocytic events^28–30^. The putative role of larger clathrin arrays in cell adhesion and migration has long been discussed^31,32^. It has recently been readdressed with the observation that specialized clathrin arrays named tubular clathrin/AP2 lattices (TCALs) are responsible for binding collagen fibres in a 3D-environment^19^ and that association of clathrin-coated plaques with the ECM is integrin dependent^27,33,34^.

To date, the mechanisms that lead to clathrin-coated plaque formation and stabilization at the plasma membrane and the cellular and extracellular determinants that dictate whether a clathrin-coated plaque displays an endocytic or a non-endocytic function are still unclear. Similarly, the molecular and/or physical determinants that drive clathrin-coated plaque formation remain poorly understood. There is evidence that some clathrin-coated plaques are resulting from frustrated endocytic events^33^. However, it remains unclear whether cells can generate specific cellular and/or extracellular signals that lead to a local induction of clathrin-coated plaques which in turn will provide the cell with a specific function.

In this work, we demonstrate that disassembling FAs, at the leading edge of the cell, are replaced by clathrin-coated plaques during cell migration, in an integrin-dependent process, that we term “switch from FAs to clathrin-coated plaques”. We could trace the signals leading to this switch back to the digestion of the ECM and the generation of extracellular topographical cues by FAs. We identified Eps15 and Eps15R as important players in the formation of clathrin-coated plaques. Migration assays of wild type and Eps15/R depleted cells, allowed us to correlate directionality of cell migration with the formation of clathrin-coated plaques at topographical cues. This work reveals a novel function of clathrin-coated plaques and demonstrates that flat clathrin arrays act as plasma membrane-associated supramolecular complexes which sense extracellular topographical cues to influence cell behaviour.

## Results

### Dynamics and ultrastructural characterization of clathrin-coated plaques

The presence of clathrin-coated plaques at the plasma membrane has been long known^31,35^, but their function is still poorly understood. It was shown that their presence was cell type specific and that they were limited to the ventral plasma membrane^36^. This side-specific localization and the fact that clathrin-coated plaques are in close contact with the substrate^31,32^ as well as associated with the ECM receptors integrins^27,32–34^ strongly suggest a function in cell adhesion. Aiming at characterizing this function, we exploited the human glioblastoma U373 cell line previously reported to display clathrin-coated plaques^36,37^. Live fluorescence microscopy of U373 stably expressing the sigma subunit of the clathrin adaptor AP2 fused to eGFP (AP2-eGFP) showed clathrin structures with high fluorescence intensity and a long lifetime as well as non-terminated events (Supplementary Fig. 1a). In contrast, transient endocytic clathrin structures had both a lower fluorescent intensity and lifetime. Transmission electron microscopy (TEM) of metal replicas confirmed the presence of flat clathrin-coated plaques with or without budding CCPs at the rim, displaying a surface area of up to 100,000 nm^2^. In comparison, the endocytic invaginated spherical CCPs display an average projected area of 15,000 nm^2^ (Supplementary Fig. 1b-e). Correlative light and electron microscopy (CLEM) of these U373 cells expressing AP2-eGFP confirmed that the high fluorescence long-lived clathrin-coated structures (Supplementary Fig. 1a) were indeed clathrin-coated plaques (Supplementary Fig. 1f-g). As such, in this work, we will define clathrin-coated plaques as long-lived clathrin coats using live-cell fluorescence microscopy. We could confirm that these structures are exclusively found at the ventral side of cells (Supplementary Fig. 2a-b) and furthermore, show that this side specific localization of clathrin-coated plaques correlates with the interaction of the cell with the extracellular environment. Cells grown on adhesive micropatterned substrates display both clathrin-coated plaques and CCPs on the adhesive parts whereas only CCPs are observed on the non-adhesive sections of the substrate (Supplementary Fig. 2c-e). In summary, in agreement with previous studies^36,37^, our results showed that U373 cells form flat clathrin-coated plaques with a long lifetime exclusively at attached plasma membrane parts.

### Clathrin-coated plaques are formed after FA disassembly

It has been proposed that the clathrin machinery is involved in recycling of FAs^38–40^. To address whether clathrin-coated plaques might be involved in this process, FAs were labelled in U373 cells by expressing the markers of FAs, zyxin, vinculin, focal adhesion kinase (FAK) or paxillin fused to the fluorescent protein mCherry. Mature FAs and smaller focal complexes did not colocalize with clathrin structures of any size, suggesting that neither CCPs nor clathrin-coated plaques have a physical correlation with FAs (Fig. 1a). To rule out that the lack of colocalization between clathrin-coated plaques and FAs was due to the transient nature of their interaction during the disassembly process, we performed live-cell fluorescence microscopy of migrating U373 cells expressing AP2-eGFP as a marker for clathrin structures and mCherry-zyxin, as a marker for FAs. Interestingly, we observed that during cell migration, clathrin-coated plaques frequently formed at the position where FAs were disassembled (Fig. 1b and e, Supplementary Movie 1). In other words, following disassembly of FAs, clathrin structures are actively assembled at the same locations (Fig. 1b). During this switch from FAs to clathrin-coated plaques, the clathrin structures and the FA did not colocalize and were mutually exclusive overtime at a given position (Fig. 1c-d). The formation of clathrin-coated plaques was a process specific to the locations of disassembled of FAs whereas the appearance of clathrin-coated plaques at other locations was a rare event (Fig. 1e). This process was not specific to zyxin as similar mutual exclusions were observed using the FA markers vinculin, FAK, and paxillin (Supplementary Fig. 3). Importantly, this phenomenon was not cell type specific as we could observe a similar switch when using other cell types also known to display clathrin-coated plaques (HT1080 and U2OS, Supplementary Fig. 4). These results report a so far undescribed switch from FAs to clathrin-coated plaques and suggest that clathrin-coated plaques have a molecular origin locally imprinted by FAs.

**Figure 1:**
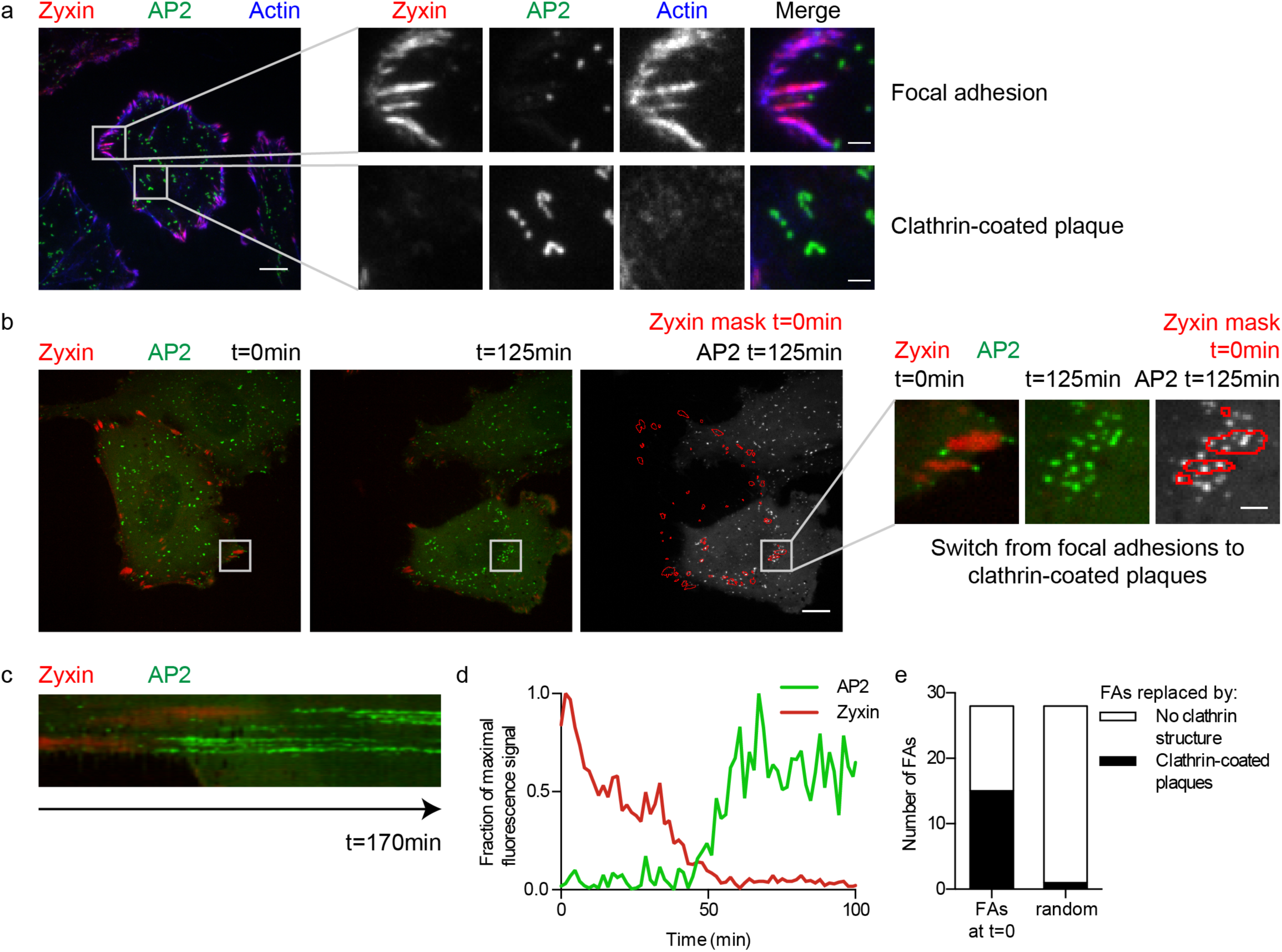
Switch from FAs to clathrin-coated plaques. (a) Representative image from total internal reflection fluorescence (TIRF) microscopy of U373 stably expressing AP2-eGFP (green) and transiently expressing mCherry-zyxin (red) stained for actin with Alexa Fluor 647-labelled phalloidin (blue). Right: Zoom on FAs labelled by mCherry-zyxin (upper panel) and on clathrin-coated plaques labelled by AP2-eGFP (lower panel). (b) Live-cell confocal spinning disc microscopy of U373 stably expressing AP2-eGFP (green) and transiently expressing mCherry-zyxin (red). Overview of a representative migrating cell at time point 0 (left) and 125 minutes later (middle). (Right) Merged images of a mask marking the mCherry-zyxin objects at time point 0 (red) and the AP2-eGFP signal 125 minutes later (green). Right: Zoom on FAs that switch to clathrin-coated plaques. (c) Kymograph of the switch from FAs to clathrin-coated plaques shown in b over 170 minutes. (d) Normalized fluorescence intensity profiles of AP2-eGFP (green) and mCherry-zyxin (red) of the switch from FAs to clathrin-coated plaques shown in b. (e) Quantification of the number of FAs being replaced by clathrin-coated plaques. Data are from live experiment performed in b. FAs at time point 0 were followed over 2 hours to quantify the number of FA replaced by clathrin-coated plaques (black) vs. the number of FA not replaced by plaques (no clathrin structures (white)). As a control the appearance of clathrin-coated plaques at randomly placed ROIs of the same size was analysed. Scale bars: 10 μm (overviews) or 2 μm (zooms).

### Clathrin-coated plaques are stabilized by integrins

In the context of integrin internalization during FA recycling, the above-described switch from FAs to clathrin-coated plaques could generate an endocytic hot spot for the internalization of integrins during FA disassembly. Most importantly, very recent reports have demonstrated the importance of integrins in stabilizing clathrin-coated plaques^33 34^. By performing immunostaining of endogenous integrins, we could show that integrins αvβ5 and β1 were enriched at clathrin-coated plaques in U373 cells (Fig. 2a-b and Supplementary Fig. 5a, respectively). This suggests that integrins are left behind at the position where FAs are disassembled and could recruit or stabilize the clathrin machinery to locally promote the formation of long-lived clathrin-coated plaques. To challenge the possible function of integrins in stabilizing clathrin-coated plaques that are formed after FA disassembly, we used the cyclic RGD peptide Cilengitide, which acts as an integrin antagonist and inhibits integrin binding to their ECM ligands^41,42^. We observed that Cilengitide treatment induced the rapid dissociation of clathrin-coated plaques (Fig. 2c-e, Supplementary Movie 2). During Cilengitide treatment, clathrin-coated plaques disassembled by the formation of several smaller transient structures resembling endocytic hot spots (Fig. 2d and Supplementary Movie 2). Together these results indicate that clathrin-coated plaques replace disassembling FAs and are recruited and/or stabilized by integrins.

**Figure 2:**
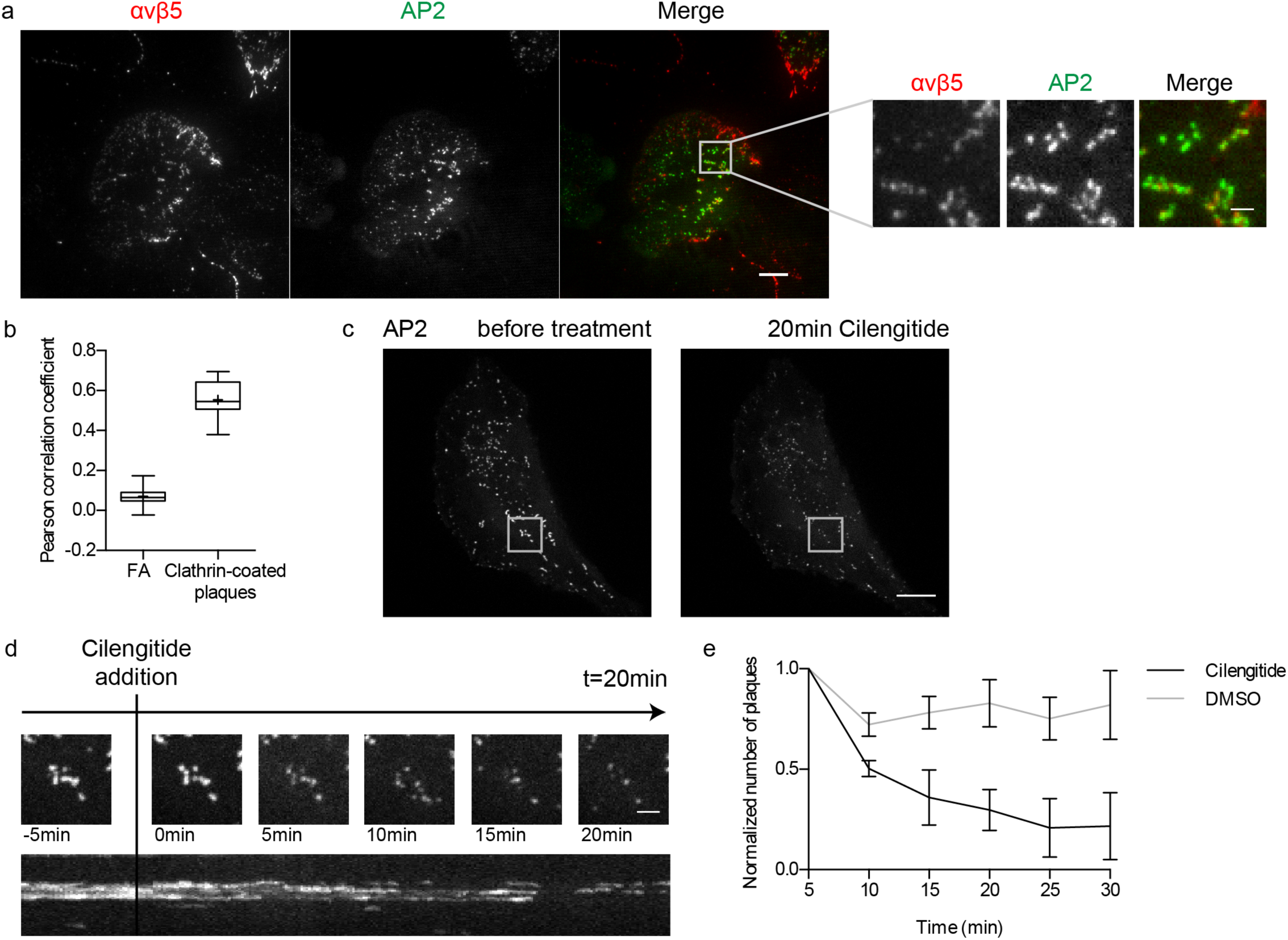
Clathrin-coated plaques are stabilized by integrins. (a) Representative image from TIRF microscopy of U373 stably expressing AP2-eGFP (green) immunostained for integrin αvβ5 (red). Right: Zoom on clathrin-coated plaques. (b) Quantification of the colocalization between αvβ5 integrin and AP2-eGFP. The graph shows the distribution of the Pearson correlation coefficient between AP2-eGFP and integrin signals at either FAs or clathrin-coated plaques. Whiskers represent 10–90 percentile, box represents second and third quartile, line marks the median and cross the mean. Results are computed from ten cells. (c) Live-cell spinning disc confocal microscopy of U373 stably expressing AP2-eGFP treated with Cilengitide (10 μM) for 20 minutes. Shown is a representative cell before (left) and after 20 minutes of treatment with Cilengitide (right). Clathrin-coated plaques were identified by monitoring the dynamic of the clathrin structures before treatment with Cilengitide. (d) Serial view of a clathrin-coated plaque (ROI shown in b) (top panel) and corresponding kymograph (bottom). (e) Quantification of the number of clathrin-coated plaques during Cilengitide (black) and DMSO (grey) treatment. Number of plaques was normalized to the time at the beginning of the treatment. Shown are the mean and SD calculated from four cells of each condition. Scale bars: 10 μm (overviews) or 2 μm (zooms).

### Extracellular signals drive the formation of clathrin-coated plaques

To address whether a specific extracellular signal created by FAs was left after FA disassembly and was in turn responsible for the integrin-dependent recruitment of clathrin-coated plaques, we monitored the switch from FAs to clathrin-coated plaques in migrating cells. We found that the location of clathrin-coated plaques is somehow defined by the extracellular environment. Indeed, we observed that clathrin-coated plaques often appear at the same locations when the same migrating cell revisits a position twice (Fig. 3a, first cell at t=0 and first cell at t=4.4h). More strikingly, if a different cell migrates over these plaque-forming areas, clathrin-coated plaques are also observed in the new cell at the same positions (Fig. 3a, merge between second cell t=7.7 h and first cell t=0 and t=4.4 h). To unambiguously demonstrate that an extracellular signal was responsible for clathrin-coated plaque formation at the same location, we correlated the location of clathrin-coated plaques on gridded coverslips when cells were seeded sequentially in multiple rounds (Fig. 3b). In this experiment, we seeded first U373 expressing AP2-eGFP on gridded coverslips and imaged clathrin dynamics to identify the locations of clathrin-coated plaques. Following EDTA-mediated removal of the first cells, a second seeding of cells was performed on the same grid and both clathrin dynamics and plaque locations were again recorded. Results show that clathrin-coated plaques formed at the same position in the first and the second round of cell seeding (Fig 3c-d). These results demonstrate that specific extracellular signals are responsible for the local induction of clathrin-coated plaques. Trypsin and KOH-based cleaning of the gridded coverslips between both rounds of cell seeding resulted in a strong reduction of the number of clathrin-coated plaques forming at the same position (Fig. 3d), suggesting that these extracellular signals are of proteinaceous nature

**Figure 3:**
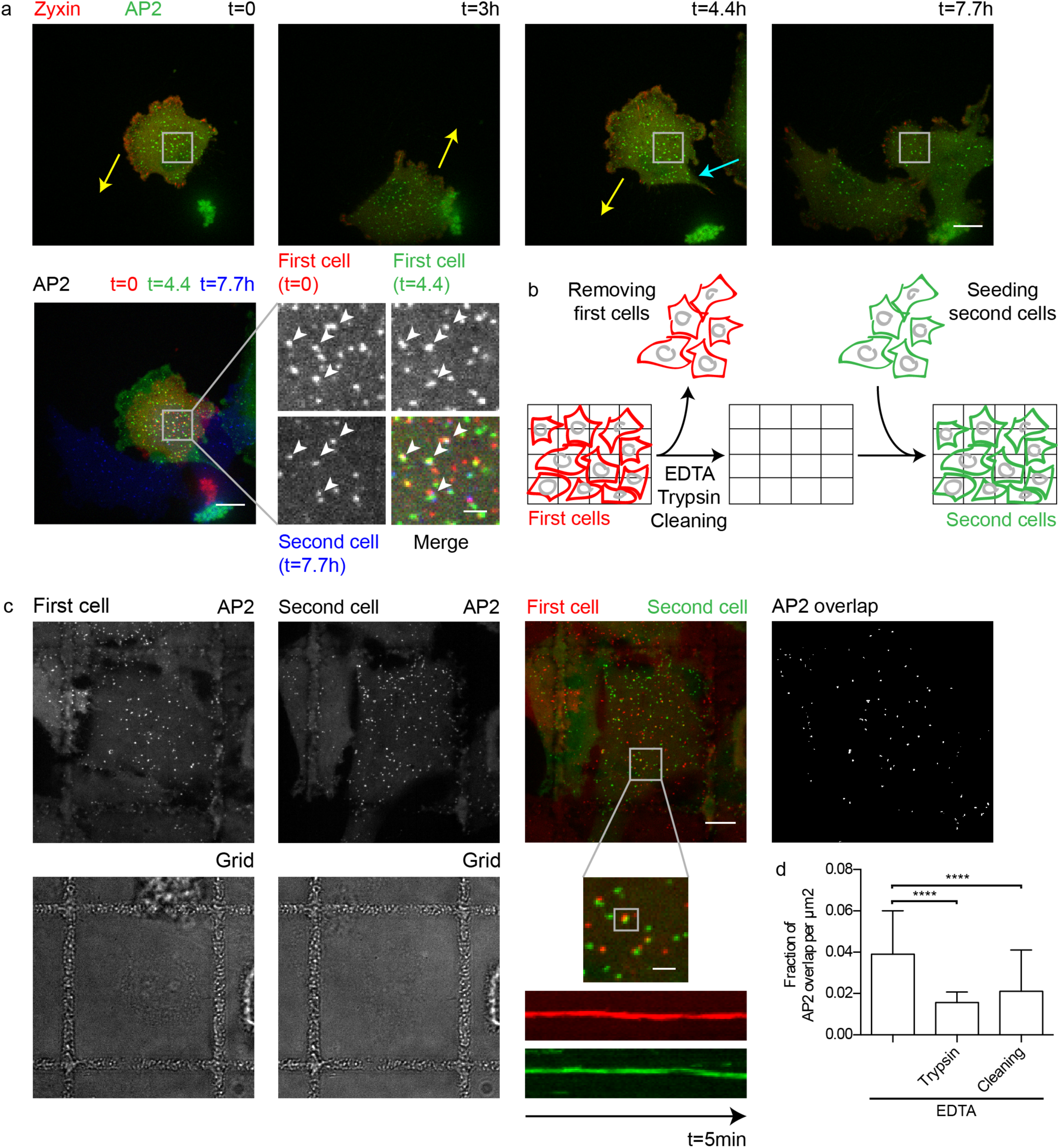
Extracellular signal induces clathrin-coated plaque formation. (a) Live-cell confocal spinning disc microscopy of U373 stably expressing AP2-eGFP (green) and transiently expressing mCherry-zyxin (red). (Top) Snapshots of a time series of two representative cells moving over the same area. Arrows point in the direction of cell migration (first cell: yellow, second cell: blue). (Bottom left) Merged images of the AP2-eGFP signal at three time points (first cell at t=0: red, first cell at t=4.4h: green, second cell at t=7.7 h: blue). Zoom on clathrin coats found at the same positions in all three time points marked by arrow heads. (b) Schematic illustrating the sequence of the experiment using gridded coverslips. A first round of cells (red) was imaged to identify the position of clathrin-coated plaques and cells were then removed by EDTA solution. The gridded coverslip was either left untreated or treated with trypsin or cleaned with a sequence of KOH, acetone, and ethanol to remove organic material. Afterwards a second round of cells (green) was seeded on the same gridded coverslip and imaged to identify the position of clathrin-coated plaques and to compare it with their position identified within the first round of seeded cells. (c) Live-cell spinning disc confocal microscopy of two rounds of U373 stably expressing AP2-eGFP on the same position of a gridded coverslip with no treatment before seeding the second round of cells. AP2-eGFP signal of first (top left) and second (top middle left) round of cells. (Top middle right) Merged image of AP2-eGFP in first cell (red) and second cell (green). Zoom in of representative overlapping clathrin coats from both rounds of seeded cells and corresponding kymographs of 5 minute long movies of clathrin-coated plaques from both rounds of seeded cells. (Top right) Binary mask of AP2-eGFP overlap. (d) Quantification of overlapping clathrin coats in the first and second round of cells using gridded coverslips. Shown are the mean with SD. Statistical analysis: t test, number of analysed images: n=31 (untreated), n=16 (trypsin), and n=15 (cleaning), P<0.01. Scale bars: 10 μm (overviews), 2 μm (zoom).

### Remodelled ECM represents an extracellular cue that promotes clathrin-coated plaque formation

FAs are supramolecular complexes not only involved in ECM binding but also responsible for ECM remodelling^13,15^. One of the important remodelling processes driven by FAs is the enzymatic digestion of the ECM^43^. To directly correlate digestion of ECM and recruitment of clathrin-coated plaques, we employed fluorescently labelled gelatin as a substitute for ECM and monitored live the digestion of the extracellular environment together with the dynamics of clathrin-coated structures. During cell migration, we could observe that FAs located at the leading edge of the cell actively digested the fluorescent gelatin, as can be seen by the appearance of dark non-fluorescent areas (Supplementary movie 3). Interestingly, by monitoring clathrin dynamics on fluorescently labelled gelatin coatings, we unravelled that the locations of long-lived clathrin-coated plaques correlated with the locations of the digested fluorescent gelatin (Fig. 4a-d and Supplementary movie 4, black area). To temporally correlate gelatin digestions by FAs and clathrin-coated plaque formation, U373 cells expressing AP2-eGFP and mCherry-zyxin were seeded on fluorescent gelatin and the dynamics of the switch from FAs to clathrin-coated plaques was observed using live fluorescence confocal microscopy. Analyses revealed that during cell migration, FAs dynamically formed and locally digested the gelatin coat, which was followed by FA disassembly and finally clathrin-coated plaque formation (Fig. 4e-g, Supplementary Movie 3). Importantly, in line with the above results showing that two different migrating cells form their clathrin-coated plaques at similar locations (Fig. 3a), our fluorescent gelatin assay reveals that the extracellular cues that lead to clathrin-coated plaques formation correspond to spots of digested ECM (Fig. 4h, Supplementary movie 5). Together, these results strongly suggest a model where FAs digest and modify the ECM creating extracellular domains responsible for the integrin-dependent recruitment of clathrin-coated plaques.

**Figure 4:**
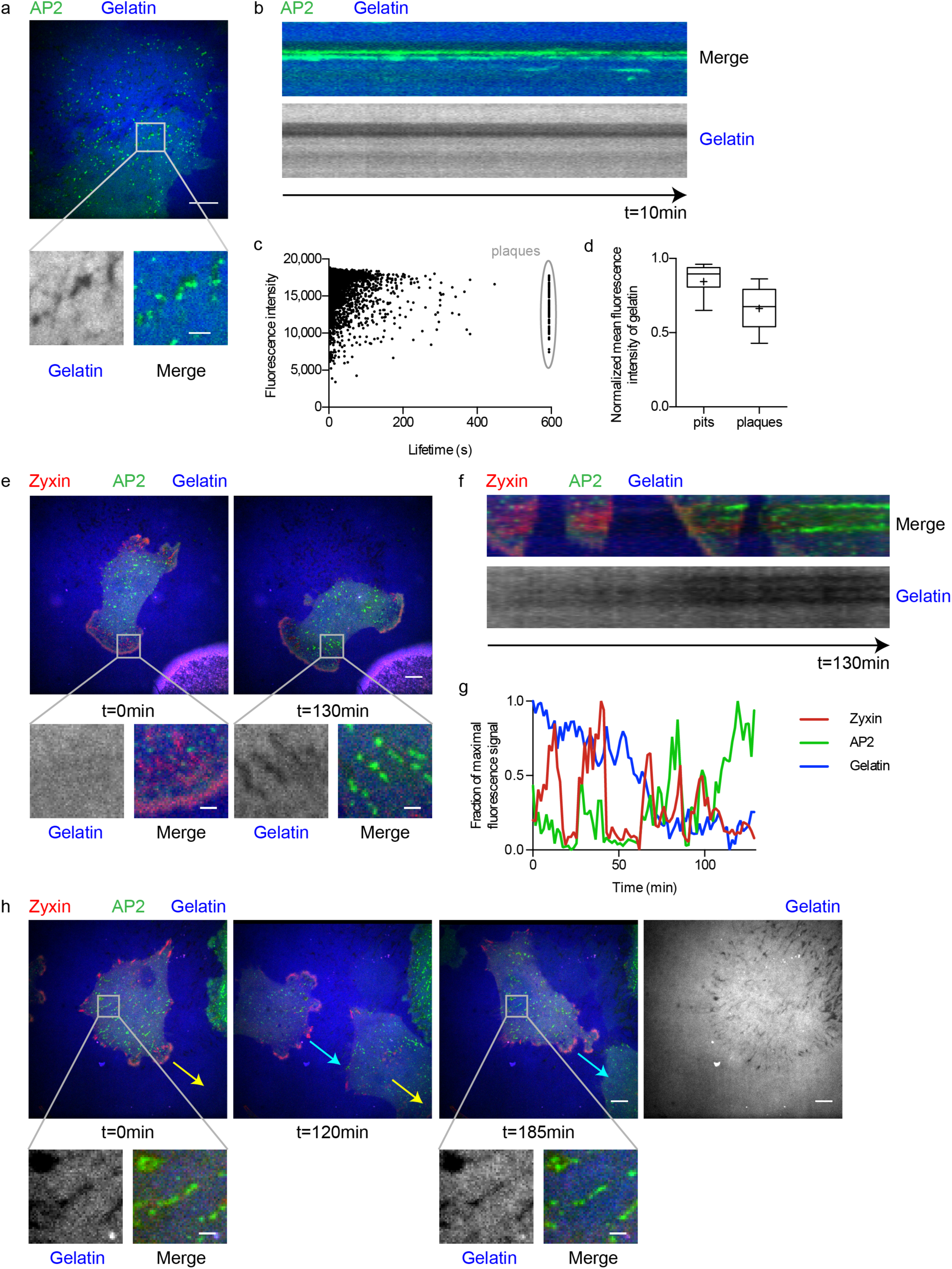
Remodelled ECM as signal for clathrin-coated plaque formation. (a) Live-cell confocal spinning disc microscopy of U373 stably expressing AP2-eGFP (green) on coverslips coated with Alexa Fluor 647-labelled gelatin (blue). Right: Zoom in of clathrin-coated plaques at digested gelatin spots. (b) Kymograph of clathrin-coated plaques and gelatin digestion shown in an over 10 minutes. (c) Lifetime vs. mean fluorescence intensity of clathrin tracks in U373 cells seeded on Alexa Fluor 647-labelled gelatin. Clathrin-coated plaques are marked by a grey circle. Shown is a plot of a representative cell. Each dot represents one CME track from a 10 minute long movie. (d) Distribution of mean fluorescence intensity of Alexa Fluor 647-labelled gelatin at transient pits and at long-lived clathrin-coated plaques (>10min) shown in c. Whiskers represent 10–90 percentile, box represents second and third quartile, line marks the median and cross the mean. Results are computed from 3,731 pits and 75 plaques. (e) Long-term live-cell confocal spinning disc microscopy of U373 stably expressing AP2-eGFP (green) and transiently expressing mCherry-zyxin (red) on coverslips coated with Alexa Fluor 647-labelled gelatin (blue). Overview of a representative migrating cell at time point 0 (left) and 130 minutes later (right). Bottom: Zoom in of FAs that switch to clathrin-coated plaques with the corresponding gelatin signal (grey). (f) Kymograph of the switch from FAs to clathrin-coated plaques on coverslips coated with Alexa Fluor 647-labelled gelatin shown in c over 130 minutes. (g) Normalized fluorescence intensity profiles of AP2-eGFP (green), mCherry-zyxin (red) and Alexa Fluor 647-labelled gelatin (blue) of the switch from FAs to clathrin-coated plaques shown in c. (h) Two representative cells migrating over gelatin digestions. Shown at time point 0 (left) and later time points (120 minutes (middle) and 185 minutes (right)). Arrows point in the direction of migration (first cell: yellow; second cell: turquois). Right: Overview of the Alexa Fluor 647-labelled gelatin signal (grey). Bottom: Zoom in time points 0 and 185 minutes showing clathrin-coated plaques at the same gelatin digestions in both cells. Scale bars: 10 μm (overviews) or 2 μm (zooms).

### Pharmacologically induced FA disassembly reveals that digestion of the ECM directly correlates with clathrin-coated plaque recruitment

To study the switch from FAs to clathrin-coated plaques and its link to FA disassembly and/or creation of extracellular cues, we exploited a pharmacological approach to induce FA disassembly. We used the ROCK inhibitor Y-27632 previously shown to induce FA disassembly^44,45^. Treatment of U373 cells with Y-27632 induced the disassembly of actin stress fibres and FAs within 20 minutes (Supplementary Fig. 6a-c). By monitoring the dynamics of both the clathrin machinery and of FAs during drug treatment, we could demonstrate that FAs are disassembled and are efficiently replaced by clathrin-coated plaques during the drug-induced FA disassembly (Fig 5a-b, Supplementary Movie 6). This replacement was similar to the process seen during cell migration as FA and clathrin-coated structures were located at the same site but temporally mutually exclusive (Fig. 5b, Supplementary Fig. 6g). Lifetime analyses of the newly formed clathrin-coated structures revealed that they were clathrin-coated plaques (Fig. 5c). Quantification of the efficiency of the drug-induced switch from FAs to clathrin-coated plaques revealed that approximately 40% of all FAs were replaced by clathrin-coated plaques (Fig. 5d). Similar to Y-27632-induced FA disassembly, treatment of cells with the myosin-II inhibitor Blebbistatin also induced FAs to switch to clathrin-coated plaques (Supplementary Fig. 6d-f and h, Supplementary Movie 7). By performing immunostaining of integrins in cells treated with Y-27632, we observed that integrins were enriched at clathrin-coated plaques located at the cell periphery and the overall colocalization of integrins with clathrin structures increased (Fig. 5e-f and Supplementary Fig. 5b). These structures corresponded to the FAs which disassembled during drug treatment, leaving behind integrins which in turn promoted the recruitment of the clathrin coat. By performing the drug-induced FA disassembly assay on labelled gelatin coatings, we could show that clathrin-coated plaques were recruited to former FA sites, which actively digested the ECM (Fig. 5g). Quantification of the frequency of drug-induced switch from FAs to clathrin-coated plaques revealed that most of the FAs which at the time of Y-27632 treatment had actively digested the fluorescent gelatin were replaced by clathrin-coated plaques, in comparison to FAs without visible gelatin digestion (Fig. 5h). These results show that in this experimental set-up, gelatin digestion is an efficient signal for clathrin-coated plaque recruitment after FA disassembly. To further correlate recruitment of clathrin-coated plaques with ECM digestion, we knocked down or overexpressed the matrix metalloproteinase MMP14 in U373 cells (Supplementary Fig. 7). MMP14 is a transmembrane metalloproteinase that plays a key role in digesting the ECM and activating other metalloproteinase and thereby regulates FAs turnover and subsequently cell migration^43,46–50^. Upon MMP14 knock-down, we observed that digestion of the fluorescent gelatin was significantly reduced (Fig. 5i-j, Supplementary Fig. 7) and this was associated with a significant reduction of the number of FAs switching to clathrin-coated plaques (Fig. 5k). Interestingly, overexpression of MMP14, which led to global digestion of gelatin coats instead of locally defined digestions as seen for WT cells (Fig. 5i-j, Supplementary Fig. 7), is also associated with a strong reduction of the drug-induced switch (Fig. 5k). Altogether, these results demonstrate that the switch from FAs to clathrin-coated plaques is tightly associated with the FA-mediated digestion of the ECM.

**Figure 5:**
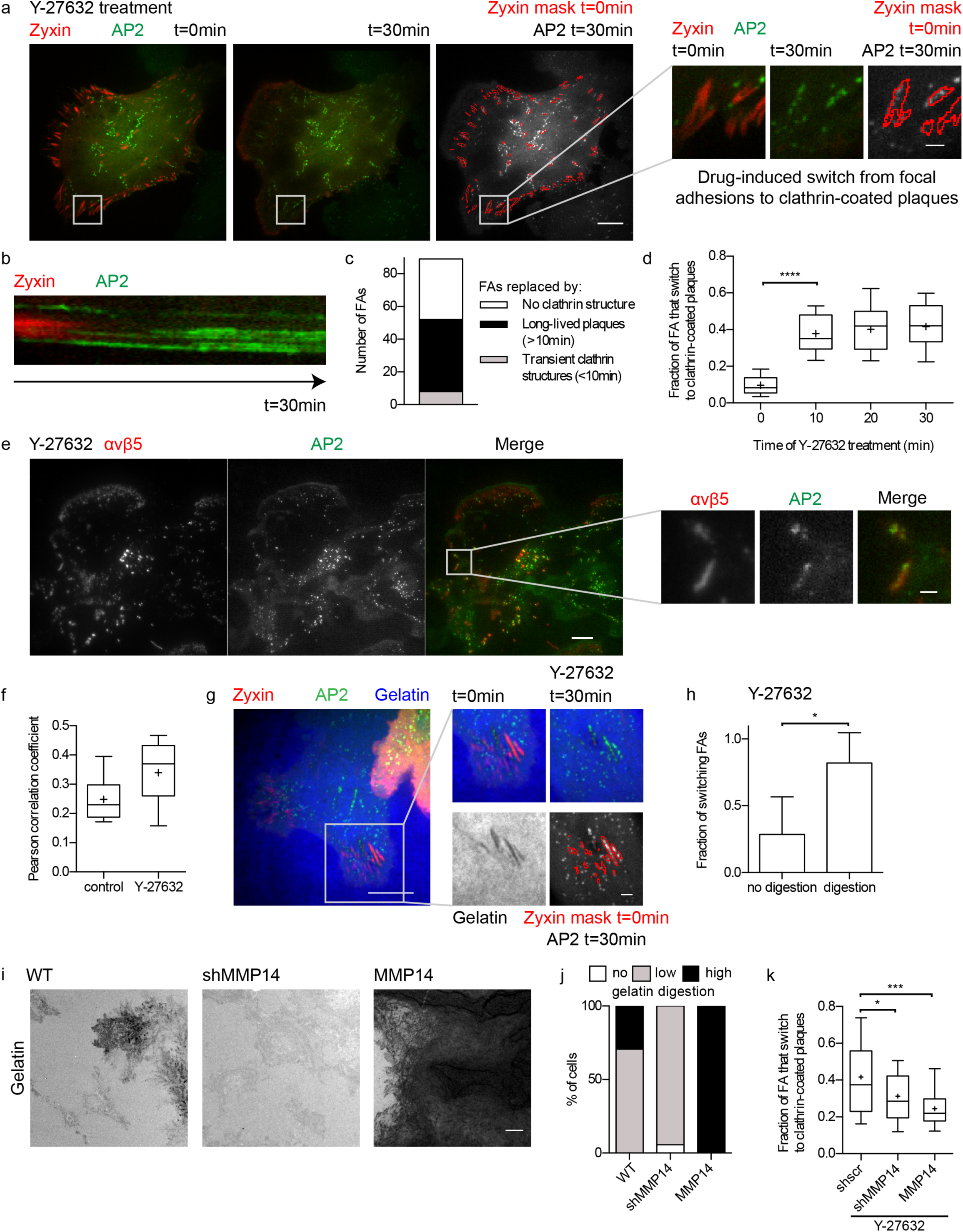
Local formation of clathrin-coated plaques by drug-induced FA disassembly. (a) Live-cell confocal spinning disc microscopy of representative U373 cell stably expressing AP2-eGFP (green) and transiently expressing mCherry-zyxin (red) treated with Y-27632 (10 μM). Cell before (left) and 30 minutes after drug treatment (middle). (Right) Merged images of a mask marking the mCherry-zyxin objects before (red) and the AP2-eGFP signal after Y-27632 treatment (green). Right: Zoom in of FAs that switch to clathrin-coated plaques during the treatment. (b) Kymograph of the switch from FAs to clathrin-coated plaques shown in b over 30 minutes of Y-27632 treatment. (c) Quantification of FA replacement by clathrin structures during Y-27632 treatment shown in a. Shown are the numbers of FAs that did not get replaced by clathrin structures (white), that got replaced by transient (<10min) clathrin structures (grey) or long-lived (>10min) clathrin-coated plaques (black). (d) Quantification of the switch from FAs to clathrin-coated plaques during Y-27632 treatment. Whiskers represent 10–90 percentile, box represents second and third quartile, line marks the median and cross the mean. Results are computed from three repetitions. Statistical analysis: t test, n=28, P<0.01. (e) Representative image from TIRF microscopy of U373 stably expressing AP2-eGFP (green) immunostained for integrin αvβ5 (red) treated with Y-27632 for 20 minutes. Right: Zoom in of clathrin-coated plaques at the periphery. Scale bars: 10 μm (overviews) or 2 μm (zoom). (f) Quantification of the colocalization between αvβ5 integrin and AP2-eGFP. The graph shows the distribution of the Pearson correlation coefficient between AP2-eGFP and integrin signals in the whole cell either non treated (control) or treated with Y-27632 for 20 min. Whiskers represent 10–90 percentile, box represents second and third quartile, line marks the median and cross the mean. Results are computed from ten cells. (g) Live-cell confocal spinning disc microscopy of U373 cells stably expressing AP2-eGFP (green) and transiently expressing mCherry-zyxin (red) on coverslips coated with Alexa Fluor 647- labelled gelatin (grey) treated with Y-27632 (10 μM). (Left) Overview of a representative cell before treatment, (right) Zoom in of FAs that switch to clathrin-coated plaques during the treatment on gelatin digestions. Scale bar: 10 μm (overview), 2 μm (zoom). (h) Quantification of the percentage of switching FAs under Y-27632 treatment for FAs without and with gelatin digestion. Shown is the mean with SD. Statistical analysis: paired t test, n=7, P<0.05. (i) Images of widefield microscopy of coverslips coated with Alexa Fluor 647-labelled gelatin with representative digested areas generated by U373 wild-type (WT), stable knock down for MMP14 (shMMP14), or stable overexpression of MMP14 (MMP14) on (red). Scale bar: 10μm. (j) Analysis of the degree of gelatin digestion by U373 WT, shMMP14 and MMP14. (k) Quantification of the switch from FAs to clathrin-coated plaques after 20 minutes of Y-27632 treatment of U373 knock down for or overexpressing MMP14. Whiskers represent 10–90 percentile, box represents second and third quartile, line marks the median and cross the mean. Results are computed from three repetitions. Statistical analysis: t test, n=28 (shscr), n=50 (shMMP14), and n=24 (MMP14), P<0.05.

### Extracellular topographical cues induce the formation of clathrin-coated plaques

Cellular processes, e.g. protein secretion, fibrillogenesis or enzymatic digestion, which occur at cell/extracellular environment contact sites, can rearrange the ultrastructural organization of the ECM^1,13^. In this way not only the protein content of the ECM can be changed, but also its physical properties^9,10,51,52^. Therefore, the above observed correlation between the location of clathrin-coated plaques and the site of gelatin digestion could be the result of a change in the topographical organization of the ECM which in turn will induce local plaque formation. To directly investigate whether local topographical cues can induce clathrin-coated plaque formation, we generated 3D-micropatterns suitable for microscopy made of an optically clear glue (Norland Optical Adhesive (NOA)) using a soft lithography approach. These 3D-micropatterns were made from optical gratings and displayed peaks of 200 nm height with a periodicity of 2 μm (Supplementary Fig. 8a-c). Analysis of the 3D-micropatterns using atomic force microscopy (AFM) revealed that they precisely reproduced the peak height and periodicity of the optical gratings (Supplementary Fig. 8b- f). U373 cells growing on such 3D-micropatterns align both their FAs and actin stress fibres parallel to the pattern as previously reported with different cell lines (data not shown and^53–57^). Interestingly, most clathrin structures were also found aligned parallel to the grating with the same periodicity of 2 μm (Fig. 6a-c). Strikingly, analyses of the clathrin structure dynamics of cells seeded on such 3D-micropatterns revealed that the long-lived clathrin-coated plaques only formed on top of the gratings whereas transient CCPs formed independently of the topographical features (Fig. 6d and e). This local induction of clathrin-coated plaque formation by substrates displaying 3D topographical micropatterns again highlights the relation between clathrin-coated plaques and extracellular physical cues.

**Figure 6:**
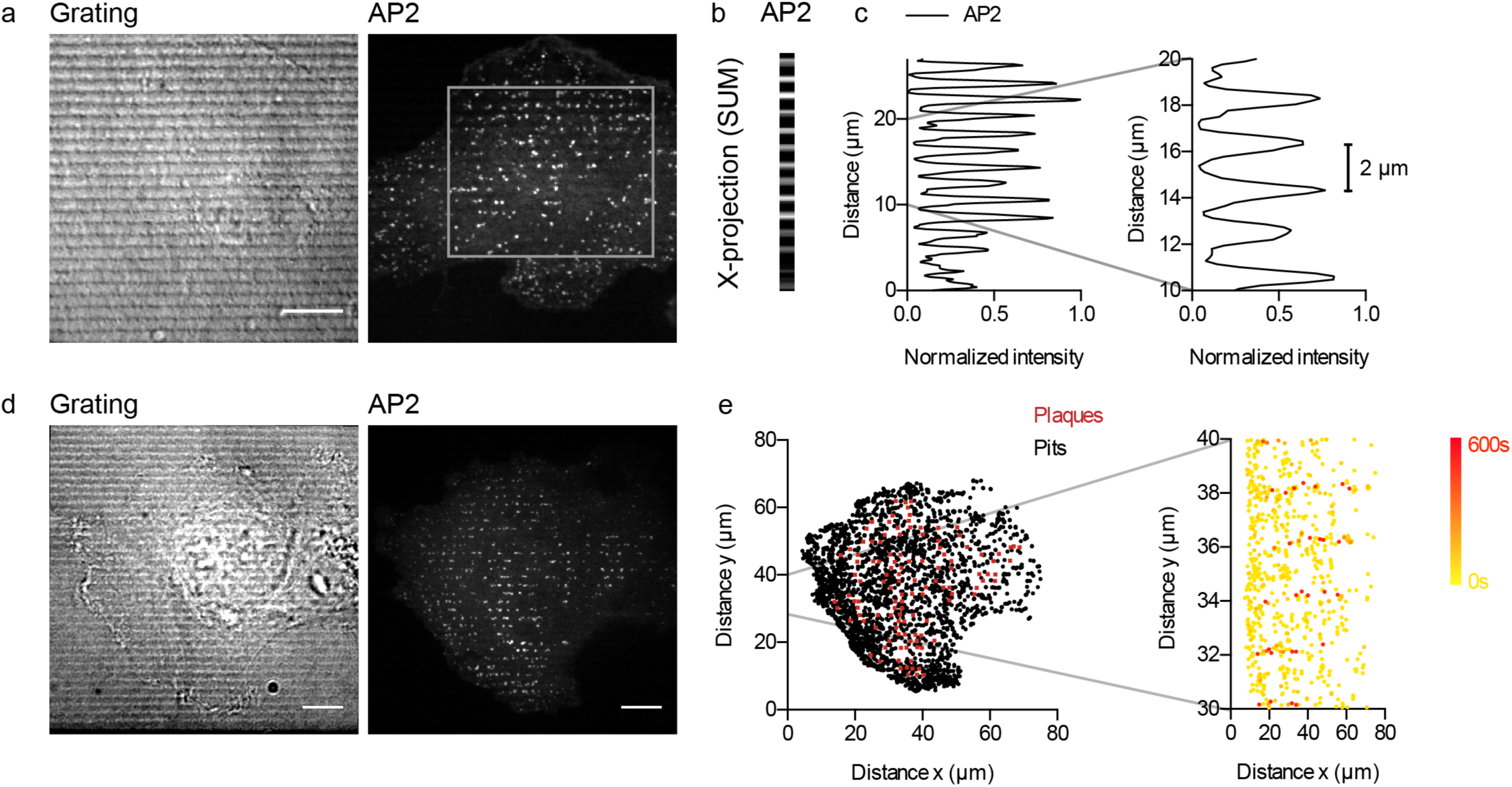
Induction of clathrin-coated plaques by topographical cues. (a) Spinning disc confocal microscopy of U373 stably expressing AP2-eGFP (right) growing on optically clear 3D-micropatterns (DIC, left). Box marks region used in b. (b) X-projection of AP2-eGFP signal. (c) Left: Normalized intensity profile from the AP2-eGFP x-projection shown in b. (d) Live-cell spinning disc confocal microscopy of U373 stably expressing AP2-eGFP (middle) growing on optically clear 3D-micropatterns (DIC, left). (e) Left: Overview of x/y-position of the tracking results from the cell shown in d. CME tracks were categories as pits (transient, black dots) and plaques (persistent for > 10 minutes, red squares). CME tracks were colour-coded according to their lifetime from yellow (0 s) to red (600 s). Scale bars: 10 μm.

### Clathrin-coated plaques are promoted by specific CME initiator proteins

Formation of clathrin-coated structures is achieved through multiple adaptor and accessory proteins, which are regulated in space and time to precisely coordinate coat assembly at the ultrastructural level^22,23^. Most studies that have looked at the functions of different proteins in clathrin coat formation have focused on their function during endocytosis. On the contrary, very little is known about proteins specifically regulating clathrin-coated plaque formation and function^27^. Previously, it was suggested that proteins of the clathrin pioneer module that initiate CME and lead to the recruitment of clathrin to the plasma membrane, e.g. FCHO1/2 and Eps15/R, stabilize the clathrin lattice early during coat formation and help to establish the hexagonal organization of flat clathrin structures^34,58–60^. Therefore, we investigated if early clathrin-related proteins of the pioneer module are important for clathrin-coated plaque formation. We found that depletion of Eps15 and Eps15R in U373 through shRNA-mediated knock-down (shEPS15/R) resulted in a dramatic reduction in the number of long-lived clathrin-coated plaques (Fig. 7a-b, Supplementary Fig. 9a). In line with their function as a pioneer module, knock-down of Eps15/R was also associated with a reduction in the number of CCP initiation events (Fig. 7a). Importantly, the switch from FAs to clathrin-coated plaques was severely impaired in cell knocked down for Eps15/R, both during cell migration (data not shown) and during Y-27632-induced FA disassembly (Fig. 7c-e, Supplementary Fig. 9b). While drug treatment was efficiently inducing disassembly of FAs, only transient CCPs formed occasionally at the sites of disassembled FAs (Fig. 7d-e). Of note, individual knock-down of Eps15 and Eps15R, also resulted in reduction of switch from FAs to clathrin-coated plaques, but to a lesser extent compared to the dual depletion of both Eps15 and Eps15R (Supplementary Fig. 9b). These results show that, in addition to their role in the initiation of CCP formation as part of their pioneer module function^60–62^, Eps15/R also play a role in regulating plaque formation and FA switch to clathrin-coated plaques.

**Figure 7:**
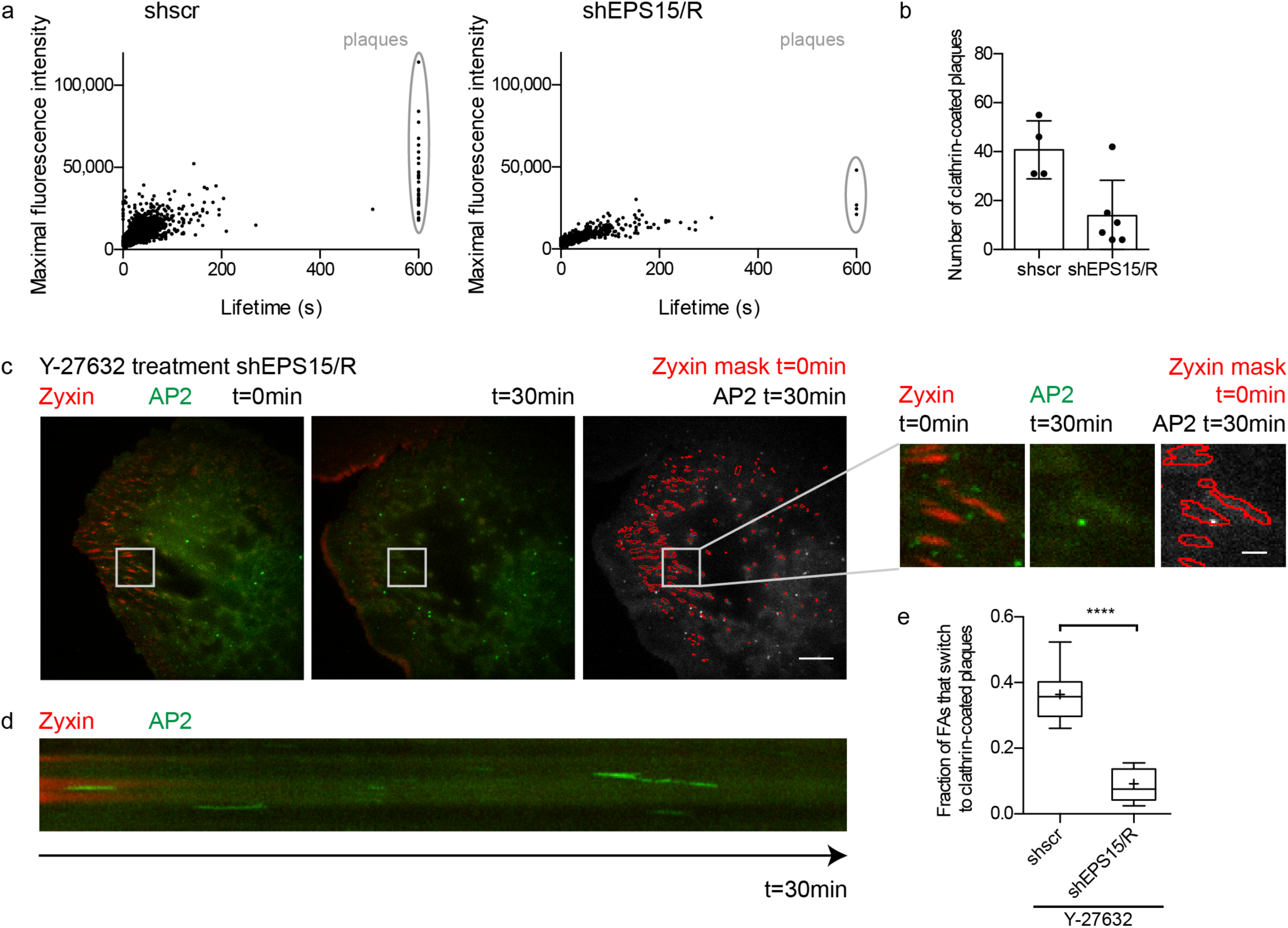
Eps15/R are regulators of clathrin-coated plaques formation. (a) Lifetime versus maximal fluorescence intensity of AP2-eGFP of CME tracks for U373 stably expressing scrambled shRNA shscr (left) and shEPS15/R (right). Persistent clathrin-coated plaques are marked by a grey circle. Shown are plots of a representative cell of each cell line. Each dot represents one tracked CME event of a 10 minute long movie. (b) Quantification of the number of clathrin-coated plaques in U373 stably expressing shscr and shEPS15/R cells. Shown is the mean with SD computed from the tracking results of at least four cells. Dots represent the results of each cell. (c) Live-cell confocal spinning disc microscopy of representative U373 shESP15/R cell stably expressing AP2-eGFP (green) and transiently expressing mCherry-zyxin (red) treated with Y-27632 (10 μM). Cell before (left) and 30 minutes after drug treatment (middle). (Right) Merged images of a mask marking the mCherry-zyxin objects before (red) and the AP2-eGFP signal after Y-27632 treatment (green). Right: Zoom on FAs that disassemble during the treatment. (d) Kymograph of the disassembly of FAs shown in b over 30 minutes of Y-27632 treatment. (e) Quantification of the switch from FAs to clathrin-coated plaques after 20 minutes of Y-27632 treatment of U373 shEPS15/R. Whiskers represent 10–90 percentile, box represents second and third quartile, line marks the median and cross the mean. Results are computed from three repetitions. Statistical analysis: t test, n=38 (shscr), n= 29 (shEPS15/R), P<0.01. Scale bars: 10 μm (overviews) or 2 μm (zooms).

### Cell migration is influenced by substrate topography through clathrin-coated plaque formation

Topographical cues of the ECM have been shown to influence the direction of cell migration^10,57,63^. This phenotype called contact guidance plays an important role during development^64–66^ as a well as tumour metastasis^67,68^. Our findings that clathrin-coated plaques can form after FA disassembly and the fact that this recruitment appears to be linked to the creation of extracellular cues in the ECM, led us to hypothesize that clathrin-coated plaques might participate in contact guidance-mediated directionality of cell migration by recognizing the extracellular cues generated by FAs in the ECM. To study the influence of clathrin-coated plaques on the directionality of cell migration, we followed U373 cells on 3D-micropatterns as well as on flat substrates made of the same material and we characterized the contact guidance-mediated directionality of cell migration. As described above, WT U373 displayed clathrin-coated plaques aligned with the 3D-micropatterns (Fig. 8a) and we found that migration of these cells was directed parallel to the grating following the grid lines (Fig. 8c, e (right panel) and g). In comparison, WT U373 cells display random cell migration on flat surfaces made of the same optically clear glue (Fig. 8e and g). As expected, Eps15/R knock-down cells seeded on the 3D-micropattern failed to display clathrin-coated plaques (Fig. 8b). Importantly, cell migration analysis of Eps15/R knock-down cells on 3D-micropatterns revealed that cells depleted of clathrin-coated plaques do not show directional migration along the grating orientation (Fig. 8b, d, f and g). Altogether, these results indicate that clathrin-coated plaques participate in contact guidance-mediated directionality of cell migration.

**Figure 8:**
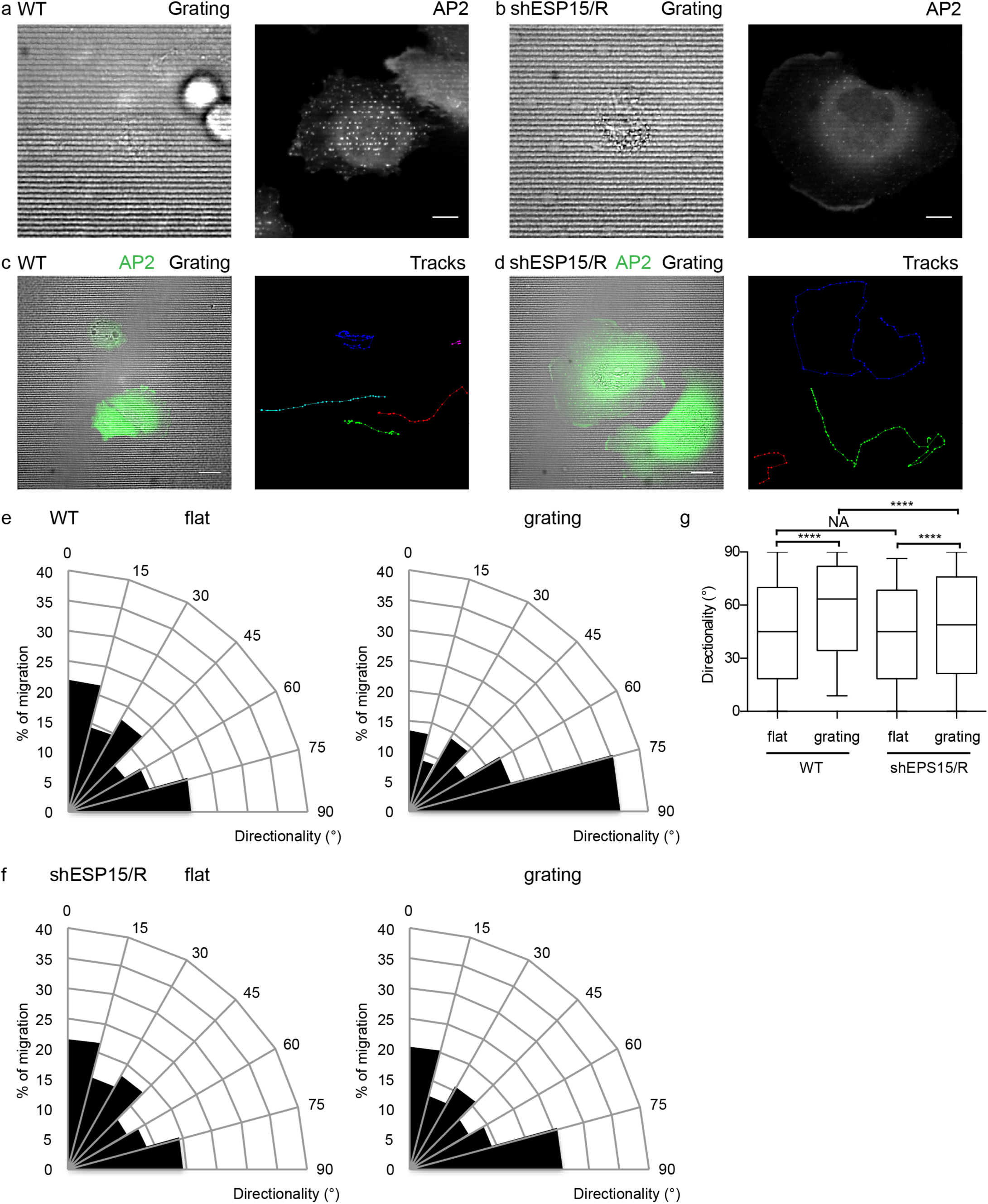
Clathrin-coated plaques establish directionality during contact guidance-mediated cell migration. U373 WT (a) and shEPS15/R (b) stably expressing AP2-eGFP seeded on optically clear 3D-micropattern. Scale bar: 10 μm. Live-cell spinning disc confocal microscopy of U373 WT (c) and shEPS15/R (d) stably expressing AP2-eGFP (green) on optically clear 3D-micropattern (grey, DIC). Migration tracks of cells are shown by multi-coloured dots and lines. Scale bar: 20 μm. Angular histogram of the directionality of migration for WT (e) and shEPS15/R (f) cells on flat substrate (left) or grating (right). (g) Distribution of directionality of cells migrating on flat substrate or 3D-micropattern. Whiskers represent 10–90 percentile, box represents second and third quartile, and the line marks the median. Statistical analysis: nonparametric t test (Mann-Whitney test), P<0.01. Directionality was calculated between time frames with a rate of 10 to 13 minutes. Results are computed from three repetitions. Number of tracks: WT: n=223 (flat), n=276 (grating), shEPS15/R: n=174 (flat), n=155 (grating).

## Discussion

Here, we describe how modifications of the ECM can lead to recruitment of clathrin-coated plaques and correlate their spatio-temporal formation with the regulation of cell migration. We demonstrate that clathrin-coated plaque formation is associated with the modification of the extracellular environment by FAs. Precisely, by combining live monitoring of ECM digestion/modifications and dynamics of clathrin-coated plaque formation, we show that, during cell migration, metalloproteinases located in FAs, digest the ECM creating specific extracellular cues which in turn drive the formation of flat clathrin arrays. Localization of clathrin-coated plaques and FAs at these specific sites reveal that these structures are temporally mutually exclusive. We name this process “FA switch to clathrin-coated plaques”. By exploiting 3D-micropatterns, we show that these extracellular signals represent topographical cues, which are recognized by the clathrin machinery inducing the local formation of plaques. We identify Eps15 and Eps15R as key regulators of plaque formation. Finally, we show that by recognizing the extracellular topographical cues, clathrin-coated plaques regulate the directionality during cell migration.

The presence of clathrin-coated plaques at the ventral plasma membrane has been long known^36^. Recently, although their origin and function remained enigmatic, several studies have reported that the presence of clathrin-coated plaques at the plasma membrane was dependent on integrins^33,34^. These findings described that not only are integrins enriched at clathrin-coated plaques, but also describes that they are mandatory to stabilize flat clathrin arrays^27,33,34^. This is consistent with our findings that interference with integrin binding to the ECM proteins impacts clathrin-coated plaque stability and induces their disassembly. The cyclic RGD peptide Cilengitide was recently shown to be a competitive antagonist which exerts its activity only on integrins which are not bound to ECM RGD motifs^69^. As such the observation that Cilengitide can induce plaque disassembly (this work and^33^) suggests that although plaques are stable structures at the plasma membrane, their integrin-dependent binding to the ECM is highly dynamic. Importantly, our work also suggests that the presence of integrin clusters is necessary, but not sufficient for clathrin-coated plaques. A great number of integrin clusters can be found at the plasma membrane, but not associated with the formation of clathrin-coated plaques (Fig. 2, Supplementary Fig. 5). Recent work by Baschieri et al. highlighted the connection of clathrin-coated plaque formation and physical properties of the substrate suggesting a mechanosensing function of plaques. They showed that stiff ECM substrates favour the formation of long-lived clathrin arrays by preventing receptors (e.g. integrins) from being internalized^33^. In light of our findings, FAs-mediated remodelling might change the physical properties of the ECM leading to the induction of clathrin-coated plaques. We could show that topographical cues of the substrate (i.e. gelatin digestions and 3D-micropatterns) are potent signals for plaque formation in an integrin-dependent manner. Furthermore, ECM remodelling can change the mechanical properties of the substrate that could be sensed by clathrin-coated plaques^70–72^.

The presence of extracellular 3D nanostructures has been previously described to be able to induce the formation of clathrin-coated structures^18–21^. Endocytically active CCPs are found to be induced both by artificial nanostructures^18,20^ and by virus particles which impose local membrane curvature^21^. Recruitment and/or induction of clathrin lattice assembly has previously been proposed to be linked with the specific recruitment of membrane curvature sensing proteins like N-BAR proteins^18^. Extracellular collagen fibres have also been reported to induce the formation of a unique clathrin coat called tubular clathrin/AP2 lattices (TCALs)^19^. Although TCALs seem to have some similarities to clathrin-coated plaques, it is not clear if they describe the same phenomenon in a different context. Both of these structures contain integrin receptors and form at locally defined ECM structures. One important difference between TCALs and clathrin-coated plaques emanating from a FA switch is that TCALs only form in a complex 3D environment and are of a more transient nature. On the contrary, clathrin-coated plaques are stable for hours and form in classical 2D cell culture but are also induced by extracellular 3D features.

We have identified Eps15 and Eps15R as key players for clathrin-coated plaque formation and for the switch from FAs clathrin-coated plaques. Additional to their function in initiation of CCPs as part of the pioneer module^60–62^, a recent study describes that together with the clathrin adaptor Numb, Eps15/R contribute to integrin β5 localization in flat clathrin lattices^34^. By binding to the integrin β5 cytoplasmic domain, the Numb-Eps15/R complex links integrins to the clathrin coat and helps to establish integrin-containing clathrin-coated plaques. This mechanism could explain why cells depleted from Eps15/R fail to form clathrin-coated plaques after FA disassembly.

During migration, we show that cells actively remodel their surrounding and generate topographical cues. Importantly, formation of these cues correlates with the integrin-dependent formation of clathrin-coated plaques after FA disassembly. Additionally, using our 3D-micropatterns, we could demonstrate that topographical cues can directly promote the formation of clathrin-coated plaques. As such, it tempting to propose that during cell migration, the FA-mediated topographical cues represent the signal leading to plaque formation. This model is supported by the fact that removing these topographical cues indeed leads to a loss of clathrin-coated plaque formation. Specifically, we could show that when cells are either depleted of or overexpressing the matrix metalloproteinase MMP14, the ECM is not digested or fully digested, respectively (Fig. 5i-j). On these surfaces which are lacking topographical cues (either because there is a lack of local digestion or because there is a complete digestion of the entire surface), switch from FAs to clathrin-coated plaques was severely impaired (Fig. 5k). Similarly, in our cell seeding experiments (Fig. 3b-c), when the ECM was removed by treatment of the surfaces using either KOH or trypsin, a loss of spatial colocalization of plaque between the two rounds of seeded cells was observed (Fig. 3d). As such we propose that FAs generate extracellular guidance patterns which will be recognized by clathrin-coated plaques ultimately influencing cell migration and establishing directionality along these topographical cues.

The exact mechanisms by which plaques can influence cell migration remains to be determined. Several studies report clustering and recruitment of transmembrane receptors and signalling molecules at clathrin-coated plaques^29–30–33^. Their stable nature would provide the cells with a long-lasting signalling platform which could influence fundamental processes regulating cell migration. However, in such a model, it is hard to explain how the presence of plaques can influence migration directionality. In the case of FA assembly and turnover, it has been proposed that internalizing integrin at the rear and their transport to the front of migrating cells might contribute to the establishment of directionality^73–74^. As such, it is possible that a similar gradient of integrin interaction with and/or integrin recycling by clathrin-coated plaques participates in directional cellular migration.

The regulation of clathrin-coated plaques and the formation of cell type-specific clathrin structures might serve as topography sensing protein complexes and influence cellular behaviour due to ECM features. In the context of a population of cells, our results support a model where a leader cell will create extracellular cues through FA switch to clathrin-coated plaques, which will then be recognized by the following cells influencing its migration. The best characterized mechanisms leading to directed cell migration is chemotaxis where cells have the ability of sensing external gradient of chemotactic factors^55^. In the complex environment of tissue in a whole organism, chemotactic gradients are further shaped by ECM (diffusion and immobilization) and neighbouring cells (secretion and sequestering)^75^. Importantly, cells not only respond to chemical gradients but also have been described to respond to mechanical gradients within the extracellular space (durotaxis)^75^. As such it is possible that clathrin-coated plaque might participate in regulating a unique type of collective migration, where cell-to-cell contact is dispensable and where the collective behaviour comes from sensing extracellular topographical cues left by the preceding cells. The formation of clathrin-coated plaques and its impact on guiding cell migration during contact guidance as well as collective cell migration might be a highly important process in development as well as tumour migration.

## Materials and Methods

### Cell culture

U373, HT1080, and U2OS cells were maintained in DMEM (Gibco, 41965-039) and Hek293T cells for Lentivirus production were maintained in IMDM (Gibco, 21980-032) both supplemented with 10% FBS (Biochrom, S0615) and Penicillin/Streptomycin (Gibco, 15140) at 37°C and 5% CO2. Sf9 were maintained in Sf-900 III SFM (Gibco, 12658019) at 27°C.

### Plasmids, shRNA and antibodies

Plasmids for mammalian expression of mCherry-FAK (55044), mCherry-paxillin (55114), and mCherry-vinculin (55160) were purchased from Addgene. Mammalian expression plasmid for mCherry-zyxin used for the cloning was a gift from Benjamin Geiger. The gateway vector pDONR 221 (12536017) and BacMam pCMV-DEST (A24223) were purchased from Invitrogen and were used for cloning of BacMam pCMV mCherry-zyxin to generate recombinant bacmid DNA for BacMam production. Mammalian expression plasmid containing the rat AP2 subunit sigma2 C-terminally fused to eGFP^76^ was used for the generation of stable cell lines. The lentiviral backbones pLKO.1 puro was a gift from Dr. Björn Tews and pLKO.1 blast (26655) was purchased from Addgene and were used for cloning of shRNA expression vectors for lentivirus production. The gateway entry vector pENTR MMP14 was obtained from the DKFZ Vector and Clone Repository and the gateway destination vector pWPI puro (Invitrogen) was used for cloning of MMP14 expression vector for lentivirus production. The VSV-G envelope expressing plasmid pMDG.2 (12259) and the lentiviral packaging plasmid psPAX (12260) were purchased from Addgene were used for lentivirus production.

The target sequences shscrambled (shscr) (5’ CAACAAGATGAAGAGCACCAA 3’), shMMP14 (5’ CATTGCATCTTCCCTAGATAG 3’), shEPS15 (5’ CCCAGAATGGATTGGAAGTTT 3’), and shEPS15R (5’ GAAGTTACCTTGAGCAATC 3’) were used for the shRNA-mediated protein knock down.

Monoclonal antibody against αvβ5 integrin (ab177004; used 1:100 for IF), MMP14 (ab51074; used 1:100 for IF and 1:5,000 for WB) and Eps15R (EP1146Y; used 1:5,000 for WB) were purchased from Abcam. Monoclonal antibody against β actin (A5441; used 1:5,000 for WB) and vinculin (V9131; used 1:800 for IF) and polyclonal antibody against Eps15 (HPA008451; used 1:5,000 for WB) were purchased from Sigma. Monoclonal antibody against β integrin (552828; used 1:1,000 for IF) was purchased from BD. The secondary antibodies anti-mouse IgG HRP (NA931; used 1:5,000 for WB) and anti-rabbit IgG HRP (NA934; used 1:5,000 for WB) were purchased from GE Healthcare. The secondary antibodies anti-mouse Alexa Fluor 568 (A-11004), anti-mouse Alexa Fluor 647 (A-21235), and anti-rat Alexa Fluor 647 (A-21247) were purchased from Invitrogen.

### Western blot (WB)

For Western blot (WB) analysis, protein samples from cell lysates were produced. Cells were detached and pelleted (300 x g, 3 minutes). After one time washing with PBS, cells were pelleted again and the cell pellet was lysed with appropriate volume of NP-40 buffer (50 mM Tris pH 8.0, 150 mM sodium chloride, 1% (v/v) NP-40, protease inhibitor (Roche, 118735800001)) for 30 minutes on ice. The cell lysate was spun down for 20 minutes at 12,000 x g and the supernatant was transferred to a fresh tube.

To measure the protein concentration of protein samples, DC (detergent compatible) Protein Assay Kit II (BioRad, #5000112) was used and compared to a bovine serum albumin (BSA) protein standard. To compare amounts of proteins of interest in different samples, the concentration of the samples was adjusted by diluting higher concentrated samples with NP-40 buffer.

Sodium dodecyl sulphate polyacrylamide gel electrophoresis (SDS-PAGE) gels were prepared with appropriate concentrations of the separating gel. Protein samples were mixed with 4x Laemmli buffer (0.2 M Tris-HCl pH 6.8, 0.05 M EDTA, 40% (v/v) glycerol, 8% (w/v) SDS, 4% (v/v) β-mercaptoethanol, 0.03% (w/v) bromophenol blue), heated for 10 minutes at 95°C and spun down for 1 minute at 12,000 × g before loading. As molecular weight marker 10 μl Precision Plus Protein Dual Color Standards (BioRad, 1610374) was separated together with the protein samples in SDS-Tris-Glycine buffer (25 mM Tris-base, 200 mM Glycine, 1% (w/v) SDS). SDS-PAGE was first performed at 80 V until samples moved through the stacking gel and then turn up to 100 V until loading dye run out of the gel. Proteins were transferred to a nitrocellulose blotting membrane (0.45 μm; GE Healthcare, #10600003) by wet blot using transfer buffer (20 mM Tri-base, 160 mM Glycine, 20% methanol) at 100 V for 1 hour. Then membranes were blocked with 5 % milk in TBST (50 mM Tris-HCl pH 7.5, 150 mM NaCl, 0.1% Tween) for 1 hour at room temperature. Incubation with primary antibody diluted in TBST with 5% milk was performed overnight at 4°C. After four washes for 5 minutes with TBST, membranes were incubated with secondary antibody diluted in TBST with 5% milk for 1 hour at room temperature. The membranes were washed four times for 5 minutes with TBST and developed with ECL Western Blotting Detection Reagents (GE Healthcare, RPN2106) or Western Bright Chemilumineszenz Substrate Sirius (Biozym, 541021) using high performance chemiluminescence films (GE Healthcare, #28906837).

### Cloning of short hairpin RNA (shRNA) encoding vector for lentivirus production

Forward and reverse oligonucleotides containing short hairpin RNA (shRNA) sequence were designed according to RNA interference (RNAi) consortium (TRC) (https://www.broadinstitute.org/scientific-community/science/projects/rnai-consortium/rnai-consortium) and ordered from Eurofins. Complementary oligonucleotides were designed to generate 5’ and 3’ overhangs after annealing complementary to AgeI and EcoRI restriction sites, respectively, for direct ligation into linearized vectors. While the EcoRI restriction site was kept intact by the oligonucleotide insertion, the AgeI restriction site was destroyed, which was used for test digestions as described later. Oligonucleotides were diluted in H_2_O at a concentration of 1 μg/μl. 2.5 μl of forward and reverse oligonucleotides were mixed with 5 μl NEB Buffer 2 (NEB, B7002S) and 40 μl H_2_O. The oligonucleotide mix was heated at 95°C for 5 minutes and slowly cooled down to room temperature for annealing of both strands. For the ligation reaction, 150 ng of pLKO.1 backbone, which has been sequentially digested with EcoRI (R0101S) and AgeI-HF (R3552S) purchased from NEB, and 2 μl annealed oligonucleotides were used. A 20 μl ligation reaction was set up using T4 Ligase (NEB, M0202S) and incubated for 1 hour at room temperature. The whole ligation mix was used for transformation into DH5a (Invitrogen, 18265017).

Colonies were tested by digestion with AgeI-HF and BamHI (NEB, R0136S). Insertion of oligonucleotides containing shRNA led to the loss of AgeI restriction site. Therefore plasmids with successful insertion showed only one linearized vector band whereas re-ligated plasmids were recognized by a double band. Plasmids were further controlled by sequencing with U6 primer.

### Lentivirus production

For lentivirus production, Hek293T cells were cultured in 10 cm dishes until 80% confluence. The cells were transfected with pMDG.2, psPAX, and the shRNA encoding pLKO.1 vector or the pWPI vector containing the gene of interest using PEI. 50 μl of PEI (1 mg/ml) was diluted in 200 μl Opti-MEM (Gibco, 31985062) and mixed well. 4 μl pMDG.2, 4 μl psPAX, and 8 μl pLKO.1 or pWPI were mixed with Opti-MEM to a final volume of 250 μl and mixed well. After 5 minutes, both solutions were mixed, vortexed and incubated for 20 minutes at room temperature. Then the transfection mix was added dropwise to cells. Culture medium was exchanged the next day. After another two days, the supernatant was harvested, centrifuged (10 minutes, 4,000 x g) and filtered through a syringe filter (Millex-HA, 0.45 μm, Millipore, SLHA033SS). For short time storage, the lentivirus containing supernatant was kept at 4°C; for long time storage, the supernatant was aliquoted and stored at -80°C.

### Gateway Cloning

The Gateway Technology (Invitrogen) was used for generating entry and expression plasmids. This highly efficient cloning system is based on site-specific recombination reactions using recombination sequences (*att*-sites) and enzymes, which catalyse recombination reactions (Clonases). This system uses a set of donor and destination vectors to quickly move DNA sequences between multiple vectors. Donor and destination vectors contain a cassette flanked by *att-*sites, which hold the *ccd*B gene for negative selection after recombination reaction and a chloramphenicol resistance gene (CM^R^) for counterselection during propagation. Therefore, donor and destination vectors need to be propagated in *E.coli* strains resistant to CcdB effects. This cassette is removed by the recombination reactions and replaced by the gene of interest. More details on the system and the procedure can be found in the manufacturer’s manual.

To generate entry vectors Gateway BP Clonase II Enzyme Mix (Invitrogen, 11789) was used. In brief, BP-recombination reactions between PCR products containing *att*B-sites and pDONR 221 containing attP-sites were set up by mixing 150 ng of PCR product with 150 ng of pDONR 221. TE buffer (10 mM Tris-HCl pH 7.4, 1 mM EDTA) was added to reach a final volume of 8 μl. Then, 2 μl of BP Clonase II Enzyme Mix were added and incubated for 1-3 hours at 25°C. 1 μl of Proteinase K (Invitrogen, AM2548) was added to the reaction mix and incubated for 10 minutes at 37°C. The reaction mix was transformed into DH5α. Plasmid DNA was isolated and sequenced with M13 forward and reverse primers.

To generate expression vectors Gateway LR Clonase II Enzyme Mix (Invitrogen, 11791) was used. In brief, LR-recombination reactions between Gateway entry vectors containing *att*L-sites and Gateway destination plasmids containing *att*R-sites were set up by mixing 150 ng of the entry vector with 150 ng of the destination vector. TE buffer was added to reach a final volume of 8 μl. Then, 2 μl of LR Clonase II Enzyme Mix were added and incubated for 1-3 hours at 25°C. 1 μl of Proteinase K was added to the reaction mix and incubated for 10 minutes at 37°C. The reaction mix was transformed into DH5a.

### BacMam production

ViraPower BacMam Expression System (Life Technologies, A34227) was used to design and produce BacMam mCherry-zyxin for transduction of mammalian cell lines following the user guide.

In brief, the Gateway cloning system was used to generate the Gateway expression plasmid BacMam pCMV mCherry-zyxin. First, a Gateway entry plasmid containing the mCherry-zyxin sequence (pENTR mCherry-zyxin) was generated. The sequence was amplified form the mammalian expression plasmid mCherry-zyxin using the primers mCherry-zyxin forward (5’ GGGGACAAGTTTGTACAAAAAAGCAGGCTCAACCATGGTGAGCAAGGGCGAGGAGGATA 3’) and reverse (5’ GGGGACCACTTTGTACAAGAAAGCTGGGTCTTACGTCTGGGCTCTAGCAGTGTG 3’) to flank the sequence with attB-sites by PCR. The PCR product was used to generate the pENTR mCherry-zyxin by BP-recombination reaction. pENTR mCherry-zyxin was then used to generate the BacMAM pCMV-DEST mCherry-zyxin expression plasmid by LR-recombination reaction.

For transposition of the mCherry-zyxin gene into the bacmid, DH10Bac (Invitrogen, 10361012) were transformed and incubated for two days at 37°C. Using a blue/white screening, positive colonies were selected which contained the recombinant bacmid. The bacmid DNA was isolated following the protocol described in the user guide.

To produce BacMam mCherry-zyxin, insect cells were transfected with the bacmid mCherry-zyxin using Cellfectin II reagent (Invitrogen, 10362100) to generate a P1 viral stock. This P1 baculoviral stock was used to amplify BacMam mCherry-zyxin by infection of insect cells. This time, the infected insect cells were culture in a spinner flask to produce a high viral titre. Two rounds of amplification were performed to produce a P2 and a P3 baculoviral stock. 3% FBS was added to the P3 baculoviral stock of BacMam mCherry-zyxin and stored in aliquots at -80°C.

### Transfection and viral transduction

Transfection of cells was done using Lipofectamine 2000 (Invitrogene, 11668027) if not stated otherwise. Cells were plated in 6-well plates one day before transfection. The next day, cells were transfected at 70-80% confluence. 2 μl DNA and 4-8 μl Lipofectamine 2000 were separately mixed with 100 μl OptiMEM. The two solutions were mixed together. After incubation for 20 minutes at room temperature, the transfection mix was added drop-wise onto the cells. For generation of stable cell lines, the growth medium was exchanged for fresh growth medium after 8 hours. The cells were put under selection two days after transfection, selected for 2 weeks and sorted using FACS. For live-cell imaging of cells transiently expressing fluorescently tagged FA proteins, the growth medium was exchange for fresh growth medium after 8 hours. The transfected cells were seeded 24 hours after transfection and imaged 6-8 hours after seeding.

For transient expression of mCherry-zyxin, cells were transduced with BacMam mCherry-zyxin. Cells were seeded with BacMam mCherry-zyxin in appropriate dishes and were used for experiments 1 day after seeding. 3.5 μl of the P3 stock was used per 10,000 cells. For lentiviral transduction for generation of stable cell lines, 10,000 cells per well were seeded into 6-well plates together with 500 μl of lentivirus containing supernatant. After 2 to 3 days, growth medium was exchange to selection medium with appropriate antibiotics. Cells were sub-cultured in selection medium for 2 weeks and then used for experiments.

### Widefield epifluorescence microscopy

If not stated otherwise, widefield epifluorescence microscopy was performed with an inverted Ti microscope (Nikon) with a 20x (0.75 numerical aperture, Plan Apo λ, Nikon) dry objective or a 40x (1.3 numerical aperture, Plan Fluor, Nikon) oil immersion objective and a digital camera (DS-Qi1MC, Nikon) equipped with LED (Lumencor-Sola) as light source.

### Spinning disc confocal microscopy

Confocal live-cell imaging was performed with an inverted spinning disc confocal microscope (Nikon, PerkinElmer) with a 60x (1.42 numerical aperture, Apo TIRF, Nikon) or 100x (1.4 numerical aperture, Plan Apo VC, Nikon) oil immersion objective and a CMOS camera (Hamamatsu Ocra Flash 4) or an EMCCD camera (Hamamatsu C9100-23B). An environment control chamber was attached to the microscope to maintain cells at 37°C and 5% CO2. If not stated otherwise, live-cell imaging was performed in DMEM without phenol red (Gibco, 21063-029) containing 10% FBS.

### Total internal reflection fluorescence (TIRF) microscopy

TIRF microscopy was performed with an inverted Ti microscope (Nikon) with objective TIRF illumination, with a 60x (1.49 numerical aperture, Apo TIRF, Nikon) oil immersion objective and an EMCCD camera (Andor iXon Ultra DU-897U).

### Quantification of FA replacement by clathrin-coated plaques during cell migration

To quantify the specific formation of clathrin-coated plaques at the position of former FAs during cell migration, live-cell confocal spinning disc was performed on U373 cells stably expressing AP2-eGFP and transiently expressing mCherry-zyxin. The position of all FAs at a given time point was used to check for replacement by clathrin-coated plaques within the following 2 hours. Only FAs that were at a position still covered by the cell after this time were analysed. Each FA was categorized into either replaced by clathrin-coated plaques or not. The formation of long-lived (>10min) clathrin-coated plaques was manually verified. As a control, random positions with the same size, which did not show any zyxin signal were analysed in the same way.

### Immunofluorescence (IF)

Cells growing on glass coverslips (#1.5, diameter 12 mm purchased from Thermo Scientific or 24 mm purchased from Marienfeld) or glassbottom dishes (ibidi, 80827) for TIRF were washed once with PBS and fixed with PFA, formaldehyde, or methanol (only for αvβ5 integrin staining). Precisely, cells were incubated either with 2% PFA or 4% formaldehyde in PBS for 20 minutes at room temperature or overnight at 4°C or with ice cooled methanol for 10 minutes at -20°C. All following steps were performed at room temperature. After three washes with PBS, cells were permeabilized with either 0.5% TritonX in PBS for 15 minutes or 0.05% saponin (only for αvβ5 integrin staining) in PBS for 10 minutes followed by a blocking step with 1% BSA in PBS for 1 hour. Samples were incubated with appropriate primary antibody dilution in 1% BSA in PBS for 1 hour. After three washes with PBS, samples were stained with appropriate secondary antibody and/or Phalloidin labelled with Alexa Fluor 647 (Invitrogen, A22287) for 45 minutes followed by four washes with PBS. For normal microscopy, samples were washed with H_2_O and mounted with ProLong Gold antifade reagent with DAPI (Invitrogen, P36931). For TIRF microscopy, samples were fixed with 4% formaldehyde solution for 20 minutes, washed three times with PBS and kept in PBS.

### Inhibitor-induced disassembly of FAs

For inhibitor-induced disassembly of FAs either the ROCK inhibitor Y-37632 dihydrochloride (Enzo, ALX-270-333) or Blebbistatin (Sigma, B0560) were used with a final concentration of 10 μM or 20 μM, respectively. The inhibitors were diluted in medium with 10% FBS. To check the effect of these inhibitors on FAs and the actin cytoskeleton, cells were incubated with pre-warmed inhibitor-containing medium and fixed with formaldehyde. FAs and the actin cytoskeleton were stained using an antibody against vinculin and phalloidin, respectively. Widefield epifluorescence microscopy was used to quantify the presence of FAs and actin stress fibres in each cell. To analyse the lifetime of clathrin structures forming after inhibitor-induced FA disassembly, each FAs visible before the application of the drug was followed over 30 minutes of treatment and categorized according to the formation of clathrin structures at its position after FA disassembly. The lifetime of clathrin structures appearing during that time were analysed manually and differentiated between transient pits (<10min) and long-lived plaques (>10min).

To quantify the inhibitor-induced switch from FAs to clathrin-coated plaques, cells expressing AP2-eGFP as well as mCherry-zyxin were used. 10 cells per sample were imaged before and after treatment with pre-warmed inhibitor-containing medium. To correct for lateral shift (pixels shifts) of the samples during the exchange of medium, the images before and after the inhibitor treatment were re-aligned using clathrin-coated plaques or other immotile fluorescent spots as reference landmarks. An automated image analysis workflow by KNIME (https://www.knime.com) was used to perform object-based colocalization analysis of FAs and clathrin structures to quantify the switch. A binary mask for the mCherry-zyxin signal before and the AP2-eGFP signal after the inhibitor treatment was used to check for overlapping objects. Only FAs that were at a position still covered by the cell after the treatment were analysed. As a control for thresholding, the number of overlapping objects of the mCherry-zyxin and the AP2-eGFP signal before the inhibitor treatment was determined and subtracted for normalization.

### Inhibition of integrins

For inhibition of integrins, the cyclic pentapeptide Cilengitide (Selleckchom, S7077) was used. The inhibitor was diluted in medium containing 10% FBS at a final concentration of 10 μM. Live-cell spinning disc confocal microscopy of CME dynamics of the same cell was performed before and after drug treatment with a frame rate of 3 seconds to follow clathrin-coated plaque disassembly. To calculate the numbers of clathrin-coated plaques, CME was tracked during 30 minutes of Cilengitide treatment. The movie was separated into 5 minute long sections and the number of clathrin structures that stayed for the whole 5 minutes was calculated for each section. This numbers were normalized to the first section directly after applying the drug.

### Quantification of colocalization between integrins and AP2

TIRF microscopy images of cells expressing AP2-eGFP and immunostained for αvβ5 were analysed using the ImageJ/Fiji plugin Coloc2. As indicated, the Pearson correlation coefficient was calculated from background corrected images either for the whole cell or areas covering FAs or clathrin-coated plaques with the same size.

### ECM degradation assay

ECM degradation assays were performed as previously described^77^. To prepare labelled gelatin, a solution of 0.2% gelatin (from porcine skin; sigma (G-2500)) in PBS was prepared. To dissolve the gelatin, the solution was heated up to 37°C for 30 minutes. For sterilization, the solution was filtered using a syringe filter membrane (Millex-GS, 0.22 μm, Millipore, SLGS033SB). To label the gelatin with Alexa Fluor 647, 500 μl of 0.2% gelatin solution was preheated to 37°C for 30 minutes. 5 μl of Alexa Fluor 647 NHS Ester (10 mg/ml, Invitrogen, A37573) was added and incubated for 1 hour at room temperature protected from light. To remove the free dye, the labelling mix was dialyzed with a Slide-A-Lyzer MINI Dialysis Device (20 kMWCO, 0.5 ml; Thermo Scientific (88402)) against PBS for 2 hours at room temperature. After changing the PBS, the mix was dialyzed overnight at 4°C. During dialysis, the labelled gelatin was protected from light.

To coat coverslips with labelled gelatin, washed coverslips were first coated with poly-L-lysine (PLL Sigma, P8920), using a 50 ng/ml solution in PBS for 20 minutes at room temperature. The coverslips were washed three times with PBS. Then the PLL coat was fixed with 0.5% glutaraldehyde (Sigma, G5882) in PBS for 15 minutes at room temperature. The coverslips were washed three times with PBS. Mixture of 0.2% labelled and unlabelled gelatin in a ratio of 8:1 was preheated to 37°C for 30 minutes. Coverslips were inverted on a drop of gelatin mix (80 μl for 12 mm diameter coverslips; 120 μl for 25 mm diameter coverslips) and incubated for 10 minutes at room temperature. Coverslips were washed three times with PBS. The coated coverslips were directly used for experiments.

To seed cells on coverslips coated with labelled gelatin, detached cells were resuspended in DMEM containing 10% FBS. The cells were pelleted by centrifugation (300 × g, 3 minutes) and washed with PBS to remove residual trypsin. Cells were pelleted again and resuspended in DMEM containing 10% FBS and plated on coverslips with fluorescent gelatin matrix (20,000 cells for 12 mm diameter coverslips; 80,000 cells for 25 mm diameter coverslips). Cells were cultured for one day on the gelatin matrix and then used for experiments.

To analyse the degree of gelatin digestion of different cell lines, widefield microscopy images were used. The labelled gelatin coat underneath each cell was manually evaluated into no, low or high digestion dependent on the loss of fluorescent signal compared to the undigested coat. For drug-induced disassembly of FAs on labelled gelatin matrix, the quantification of the switch from FAs to clathrin-coated plaques was performed as described before. Additionally, each FA was evaluated manually for gelatin digestion.

### Production of 3D-micropatterns by soft lithography

To prepare 3D-micropatterns suitable for microscopy, soft lithography with a PDMS master and optically clear glue was performed. The protocol for PDMS master production was adapted from Lücker *et al*., 2014^78^ and the protocol for 3D-micropattern production with optically clear glue from Ray *et al*., 2017^79^.

To produce the PDMS master, diffraction gratings (Edmund Optics, #54-510) with a groove density of 500 grooves per mm were glued into a 10 cm petri dish using double-faced adhesive tape. One clip-pack PDMS (SYLGARD 184, 10 g clip-pack; Sigma, #761036) was mixed, poured in the petri dish, and de-gased for 1 hour. The PDMS was cured at 50°C overnight. The PDMS on the diffraction gratings was cut into small squares (approximately 1 x 1cm) and peeled off.

These PDMS masters were used to generate 3D-micropatterns made of Norland optical adhesive (Norland Products, NOA73). A drop of NOA 73 was put on a coverslip and the PDMS master was placed on top of this drop. NOA 73 was cured by UV light (UV Stratalinker 1800, Stratagene) for 15 minutes and the PDMS master was peeled from the coverslip leaving behind the micropatterned optically clear NOA 73 film. The 3D-micropatterns were illuminated with UV light a second time for 15 minutes to ensure completeness of curing. These 3D-micropatterns were sterilized with ethanol before usage.

### Sequential seeding of cells on gridded coverslips

80,000 cells were seeded in μ-Dish 35 mm (high Grid-50 Glass Bottom; ibidi, 81148). 12 hours after seeding, cells on grids were imaged by live-cell confocal microscopy. Afterwards, cells were removed with 15 mM EDTA in PBS for 10 minutes. After the removal of cells, the gridded coverslips were either left untreated, treated with 0.05% Trypsin/EDTA for 20 minutes at 37°C or cleaned by sequential sonification in 1 M KOH, acetone, ethanol, and H_2_O for 10 minutes. Another round of cells was seeded as described above. Live-cell confocal imaging was performed 12 hours after seeding at the positions imaged in the first round of cells. To evaluate the amount of clathrin structures formed at the same position in both rounds of cells, the images were aligned with ImageJ/Fiji using the landmarks plugin. Three landmarks from the grid were selected in both pictures and the second image was registered to the first image with the Best Rigid Registration method. An automated image analysis workflow by KNIME was used to perform object-based colocalization analysis of clathrin structures for quantification. A binary mask for the AP2-eGFP signal in both images was used to check for overlapping objects and to quantify the percentage from all clathrin structures found in the first round of cells. Only clathrin structures at regions covered by cells in both images were analysed.

### Tracking of clathrin structures

For the tracking of CME events, we used ilastik (http://ilastik.org). First the images were segmented using the pixel classification and object classification workflow. For tracking, the automatic tracking workflow was used. The maximal distance was put to 5 to avoid merging of close tracks. Tracking results were further analysed by automated workflows using KNIME to calculate features (lifetime, maximal fluorescence intensity and average position) of each CME event.

### Tracking of cell migration

For the tracking of cell migration, we used the ImageJ/Fiji plugin Manual Tracking. The position of a cell was manually defined by the centre of its cell body for each time point. The tracking results were further analysed by automated workflows using KNIME to calculate the directionality of cell migration. The directionality was defined as the angle between two sequential positions of a cell.

## Supporting information

## Acknowledgments

This work was supported by research grants from the Chica and Heinz Schaller Foundation and the Deutsche Forschungsgemeinschaft (DFG) in TRR186 (project 09) to SB. DB was supported by a fellowship from the Hartmut Hoffmann-Berling International Graduate School of Molecular & Cellular Biology (HBIGS) at the Heidelberg University, by a travel collaboration grant from the Boehringer Ingelheim Fonds and from the DFG via the SFB1129 (project 14). SB is members of the cluster of excellence CellNetworks. JWT is supported by the Intramural Research Program of the US National Heart Lung and Blood Institute (NHLBI), National Institutes of Health (NIH). We would like to thank Ulrike Engel and the Nikon Imaging Center (Heidelberg University) for support with TIRF microscopy, Vibor Laketa from the Department of Infectious Diseases, Virology (University Hospital Heidelberg) for support with spinning disc microscopy, and the US National Heart Lung and Blood Institute (NHLBI) Electron Microscopy Core and Light Microscopy Core facilities for use of equipment.

## Author contributions

DB designed and performed experiments, analysed data and wrote manuscript. MM designed and established BacMam mammalian expression system. VS performed drug-induced FA disassembly assays and designed 3D-micropattern experiments. CH designed, performed and analysed AFM measurements. CZ and EACA designed adhesive micropattern experiments. KAS and JWT designed and performed TEM and CLEM experiments. SB supervised the project, designed experiments, interpreted data and wrote manuscript. The authors declare that they do not have competing financial interests.

