## Supplementary material for "Focal adhesion-generated cues in extracellular matrix regulate cell migration by local induction of clathrin-coated plaques"

### *Supplementary Movies*

**Supplementary Movie 1: Switch from FAs to clathrin-coated plaques during cell migration.** Live-cell confocal spinning disc microscopy of U373 stably expressing AP2-eGFP (green) and transiently expressing mCherry-zyxin (red). Scale bar: 10µm. Movie time is displayed in the top left corner.

**Supplementary Movie 2: Disassembly of clathrin-coated plaques by integrin inhibition.** Live-cell confocal spinning disc microscopy of U373 stably expressing AP2-eGFP treated with Cilengitide (10 µM) for 20 minutes. Scale bar: 10µm. Movie time is displayed in the top left corner.

**Supplementary Movie 3: Switch from FAs to clathrin-coated plaques at local gelatin digestions during cell migration.** Live-cell confocal spinning disc microscopy of U373 stably expressing AP2-eGFP (green) and transiently expressing mCherry-zyxin (red) on coverslips coated with Alexa Fluor 647-labelled gelatin (blue). Scale bar: 10µm. Movie time is displayed in the top left corner.

**Supplementary Movie 4: Clathrin-coated plaque formation at local gelatin digestions.** Live-cell confocal spinning disc microscopy of U373 stably expressing AP2-eGFP (green) on coverslips coated with Alexa Fluor 647-labelled gelatin (blue). Scale bar: 10µm. Movie time is displayed in the top left corner.

**Supplementary Movie 5: Formation of clathrin-coated plaques in two migrating cells at the same local gelatin digestions.** Live-cell confocal spinning disc microscopy of two U373 stably expressing AP2-eGFP (green) and transiently expressing mCherry-zyxin (red) on coverslips coated with Alexa Fluor 647-labelled gelatin (blue). Scale bar: 10μm. Movie time is displayed in the top left corner.

**Supplementary Movie 6: Switch from FAs to clathrin-coated plaques during ROCK inhibition.** Live-cell confocal spinning disc microscopy of representative U373 cell stably expressing AP2-eGFP (green) and transiently expressing mCherry-zyxin (red) treated with Y-27632 (10 μM) for 30 minutes. Scale bar: 10μm. Movie time is displayed in the top left corner.

**Supplementary Movie 7: Switch from FAs to clathrin-coated plaques during myosin-II inhibition.** Live-cell confocal spinning disc microscopy of representative U373 cell stably expressing AP2-eGFP (green) and transiently expressing mCherry-zyxin (red) treated with Blebbistatin (20 μM) for 30 minutes. Scale bar: 10μm. Movie time is displayed in the top left corner.

**Supplementary Movie 8: Clathrin-coated plaques form at topographical cues.** Spinning disc confocal microscopy of U373 stably expressing AP2-eGFP (right) growing on optically clear 3D-micropatterns (DIC, left). Scale bar: 10μm. Movie time is displayed in the top left corner.

**Supplementary Movie 9: FAs disassembly in U373 shEPS15/R during ROCK inhibition.** Live-cell confocal spinning disc microscopy of representative U373 shEPS15/R cell stably expressing AP2-eGFP (green) and transiently expressing mCherry-zyxin (red) treated with Y-27632 (10 µM) for 30 minutes. Scale bar: 10µm. Movie time is displayed in the top left corner.

### *Supplementary Materials and Methods*

#### *EM and CLEM of clathrin structures using metal replicas of unroofed membrane sheets*

EM and CLEM of clathrin structures were performed as previously described[1]. Cells were seeded on PDL-coated coverslips (25 mm). Unroofing was performed 16 hours after seeding. The cells were washed three times with stabilisation buffer (30 mM HEPES pH 7.4, 70 mM KCl, 5 mM $MgCl_2$). Unroofing was performed in 2% paraformaldehyde (PFA) in stabilisation buffer using two short sonication pulses. Samples were immediately put into fresh 2 % PFA solution and fixed for 20 minutes at room temperature.

For CLEM, widefield epifluorescence microscopy of unroofed plasma membranes from AP2-eGFP expressing cells was performed with a Nikon N-STORM microscope with a 100x oil immersion objective and an EMCCD camera (Andor Ixon Ultra DU-897). To cover an area of 1 $mm^2$ a montage of 15 x 15 images with an overlap of 15% for stitching was taken. The imaged area was marked with a circle (4 mm in diameter) around the centre of the imaged area using an objective diamond scriber. The immersion oil was carefully removed from the bottom of the glass coverslip and the sample was prepared for EM.

Coverslips with unroofed membranes were fixed with 2% glutaraldehyde in PBS overnight. Samples were incubated with 0.1% tannic acid for 20 minutes at room temperature. After four washes with water, the samples were incubated with 0.1% uranyl acetate for 20 minutes at room temperature. After two washes with water, samples were dehydrated with a series of ethanol solutions (15% - 100%). Samples were placed in each ethanol solution for 5 minutes. After replacing the 100% ethanol solution, the samples where dried in a critical point dryer. The samples were then put under vacuum until they were coated. The samples were coated with JFDV JOEL Freeze Fracture Equipment with a first layer of platinum with an angle of 17[illegible] while rotating and with a second layer of carbon with an angle of 90° while rotating. For better orientation, the marked area of the coated samples were imaged with a phase contrast microscope. The samples were then cut to fit on EM grids (TED PELLA, 75 Mesh Copper, Support Films Formvar/Carbon). 5% hydrofluoric acid was used to remove the glass from the metal replica. The floating metal replica was extensively washed with water and then carefully placed on a glow discharged EM grid.

Samples were dried on filter paper and again imaged with a phase contrast microscope. TEM imaging was performed using a JEOL 1400 equipped with SerialEM freeware for montaging. Montages of large membrane sheets at 12,000 magnification (1.82 nm per pixel) with 10% overlap were imaged.

#### *Transformation of images for CLEM*

Transformation of images for CLEM was performed as previously described[1]. The FM image and the EM montage of the same membrane sheet were first manually and roughly overlaid using Photoshop. MATLAB was used to transform the fluorescence image according to the EM montage using three manually identified clathrin-coated structures. For the transformation, the centre of the clathrin structure in the electron microscope montage and the centre of the fluorescence signal determined by a Gaussian fit were used as landmarks.

#### *TEM and CLEM analysis of clathrin coats*

Analysis of TEM and CLEM data was performed as previously described[1]. For the TEM analysis, clathrin-coated structures were manually identified. The projected area was measured by outlining their surrounding using ImageJ/Fiji (https://fiji.sc). Classification into flat, dome, pit and flat with budding dome/pit structures was done by visual evaluation of their curvature. Flat structures could be distinguished from curved structures. Within the curved coats, pit structures were discriminated from dome structures by the existence of a highly contrasted surrounding ring, which represents a constricted neck. The circularity of the outlines of clathrin-coated structures was measured using the shape descriptors tool in ImageJ/Fiji. For the CLEM analysis, the fluorescent intensity was correlated to clathrin-coated structures identified in TEM.

#### *Live-cell microscopy of CME at bottom and top plasma membrane*

For live-cell microscopy of CME at the bottom and top plasma membrane, cells stably expressing AP2-eGFP were seeded on PDL-coated coverslips. Cleaned coverslips were sterilized with ethanol and washed three times with PBS. Coverslips were inverted on a drop of PDL hydrobromide (mol wt 70,000 - 15,000, 0.01% (w/v) in $H_2O$; Sigma, P6407) and incubated for 5 minutes at room temperature followed by three washes with $H_2O$. CME dynamics at bottom and top plasma membrane of the same cell were imaged with live-cell spinning disc confocal microscopy 12 hours post-seeding. Therefore, first AP2-eGFP dynamics were monitored for 10 minutes with a frame rate of 5 seconds at the bottom plasma membrane. Afterwards CME dynamics of the top plasma membrane were imaged by taking a z-stack of five images covering 2 µm in height of the very top of the cell with a rate of 5 seconds between the z-stacks. Maximum projections of each z-stack were generated and further used for tracking of CME at the top plasma membrane.

#### *Generation of adhesive micropatterns by photo-patterning*

Adhesive micropatterns have been generated by photo-patterning as previously described[2]. In brief, an UVO-cleaner (Model 342, Jelight) was used to illuminate samples with deep UV. A synthetic quartz mask with customized features was purchased from Toppan. Coverslips were activated by illuminating with deep UV for 10 minutes. For passivation, the coverslips were inverted on a drop of 0.1 mg/ml Poly(L-lysine)-graft-poly(ethylene glycol) co-polymer (PLL-g-PEG) (SuSoS Surface Technology, PLL(20)-g[3.5]-PEG(2)) in 10 mM HEPES (pH 7.4) (50 µl for 12 mm diameter coverslips; 100 µl for 24 mm diameter coverslips) and incubated for 30 minutes. To visualize passivated areas, a 1:1 mix of unlabelled and Atto633-labelled PLL-g-PEG (SuSoS Surface Technology, PLL(20)-g[3.5]-PEG(2)-Atto633) was used. Passivated coverslips were washed with $H_2O$ for 10 minutes. Before usage the quartz mask was washed with acetone and isopropanol and dried. The quartz mask was activated by illuminating with deep UV for 5 minutes on the metal bearing side to clean the surface and make it hydrophilic. A drop of 4 µl of millipore $H_2O$ was put on the quartz mask and the coverslip was placed on the drop with the pegylated side facing down. Air bubbles were removed by pressing the coverslip against the mask. Deep UV was illuminated for 3 minutes through the mask with the quartz side facing the light source. After illumination, an excess of $H_2O$ was used to remove coverslips from the quartz mask by floating. The coverslips were washed with PBS and directly used for experiments.

#### *Surface topography measurements using AFM*

AFM measurements were performed on a MultiMode 8 (Bruker). For topography measurements, an area of 12 µm x 12 µm of the surface was scanned in peak force tapping mode with a scanasyst-air cantilever (resonant frequency 70 kHz, spring constant 0.4 N/m, tip radius 5-10 nm; Bruker). AFM images were processed using gwyddion (http://gwyddion.net). Line profiles were computed by averaging profiles of 1-3 μm along the gratings.

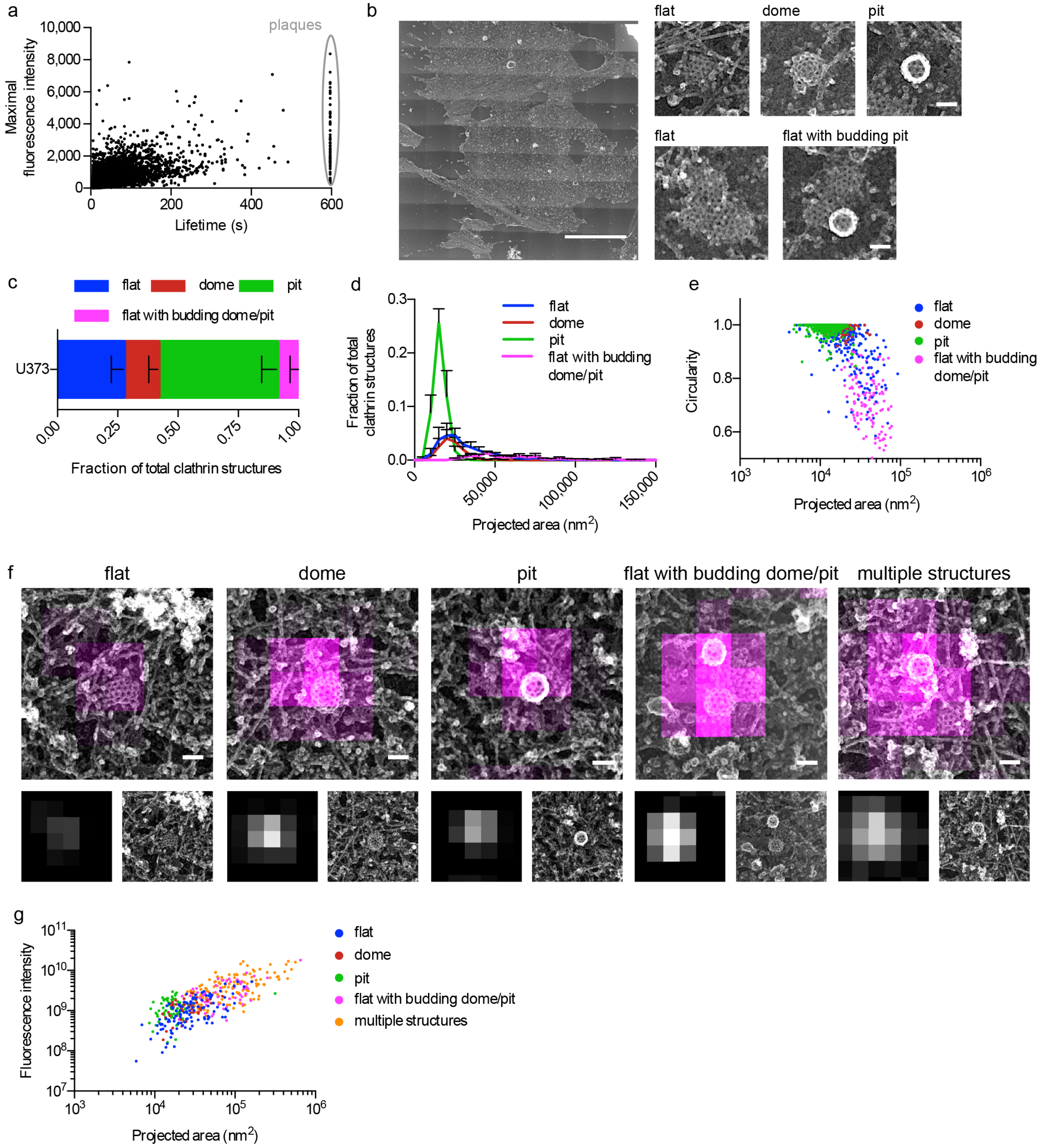
a
plaques
Maximal fluorescence intensity
10,000
8,000
6,000
4,000
2,000
0
0
200
400
600
Lifetime (s)
b
flat
dome
pit
flat
flat with budding pit
c
flat
dome
pit
flat with budding dome/pit
U373
0.00
0.25
0.50
0.75
1.00
Fraction of total clathrin structures
d
Fraction of total clathrin structures
0.3
0.2
0.1
0.0
0
50,000
100,000
150,000
Projected area (nm$^2$)
flat
dome
pit
flat with budding dome/pit
e
Circularity
1.0
0.8
0.6
$10^3$
$10^4$
$10^5$
$10^6$
Projected area (nm$^2$)
flat
dome
pit
flat with budding dome/pit
f
flat
dome
pit
flat with budding dome/pit
multiple structures
g
Fluorescence intensity
$10^{11}$
$10^{10}$
$10^9$
$10^8$
$10^7$
$10^3$
$10^4$
$10^5$
$10^6$
Projected area (nm$^2$)
flat
dome
pit
flat with budding dome/pit
multiple structures

**Supplementary Figure 1: Dynamical and morphological characterization of clathrin structures in U373 cells.**

(a) Lifetime versus maximal fluorescence intensity of AP2-eGFP of CME in U373 cells. Persistent clathrin-coated plaques are marked by a grey circle. Shown is a plot of a representative cell. Each dot represents one tracked CME event of a 10 minute long movie. (b) TEM of metal replicas of unroofed plasma membrane sheets from of U373. Left: Overview of whole membrane. Right: Zooms on representative examples of morphological categories found in the U373 cells: Flat, dome, pit, and flat with budding dome/pit. Scale bar: 10 µm (overview); 100 nm (zooms). (c) Fraction of flat (blue), dome (red), pit (green), and flat with budding dome/pit (magenta) clathrin-coated structures on plasma membrane sheets. Results are calculated from four different membranes. Total number of clathrin-coated structures: 3,627. Means with SD are shown. (d) Projected area distribution of the different clathrin morphologies found on plasma membrane sheets of U373. Means with SD are shown. (e) Circularity and size of clathrin-coated structures with different morphologies. Each dot represents one clathrin coat. Shown are the results of one representative plasma membrane sheet. (f) Examples of flat, dome, pit, flat with budding dome/pit, and multiple structures, which cannot be distinguished by FM, observed with CLEM. Top panel: CLEM, lower left: fluorescence microscopy, lower right: TEM. Scale bar: 100 nm. This panel has been adapted from Bucher *et al.*, 2018[1]. Published by Springer Nature Limited under a Creative Commons Attribution 4.0 International Licence (http://creativecommons.org/licenses/by/4.0/). (g) Correlation of size and AP2-eGFP fluorescence intensity of all clathrin structures categorized by their morphologies (flat (blue), dome (red), pit (green), flat with budding dome/pit (magenta), and multiple structures (orange)). Graph shows representative CLEM analysis of one plasma membrane sheet of U373. Total number of clathrin structures: 427.

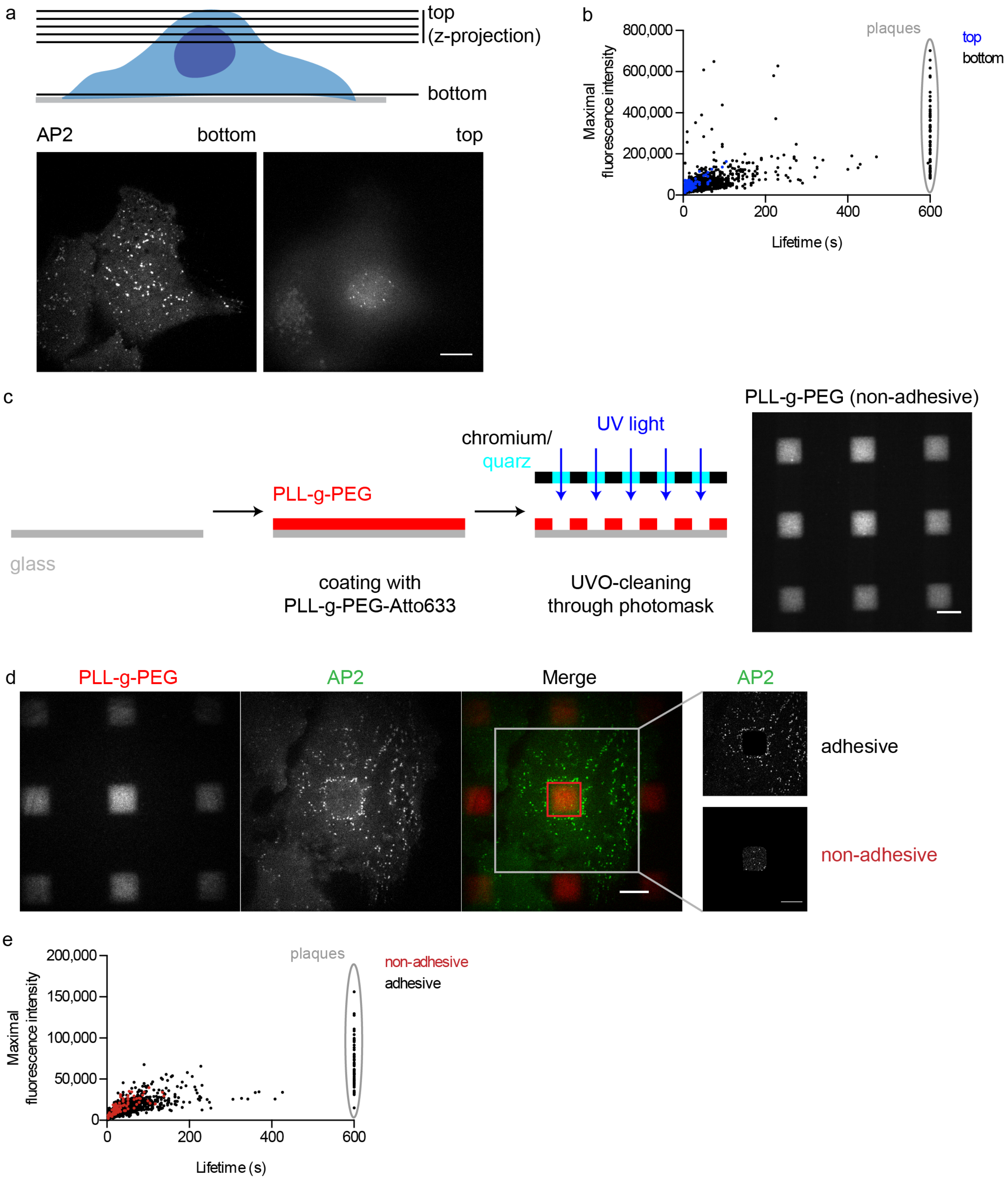


171 **Supplementary Figure 2: Clathrin-coated plaques form exclusively at attached plasma**
172 **membrane parts.**

(a) Schematic of imaging of CME at bottom and top plasma membrane. AP2-eGFP was imaged at the bottom plasma membrane in a single focal plane, for 10 minutes with a frame rate of 5 seconds. The top plasma membrane was imaged by taking a z-stack of 5 images with a spacing of 0.5 µm at the very top of the same cell. For tracking of CME on the top plasma membrane, a maximal intensity z-projection was used. Scale bar: 10 µm. (b) Lifetime versus maximal fluorescence intensity of AP2-eGFP of CME tracks from the top (blue) and the bottom (black) plasma membrane of a representative U373 cell. Each dot represents one tracked CME event of a 10 minute long movie. Persistent clathrin-coated plaques are marked by a grey circle. Number of tracks at bottom PM: 1,874. Number of tracks at top PM: 118. (c) Schematic illustrates the production of adhesive micropatterns using photo-patterning. First, glass coverslips were passivated by coating with Atto633-labelled PLL-g-PEG. Spatially controlled UVO-cleaning through a photomask was performed to generate a micropattern of adhesive and non-adhesive regions. Right: Image of the adhesive micropattern used in this experiment consisting of 10 µm x 10 µm squares of PLL-g-PEG with a spacing of 10 µm. Scale bar: 10 µm. (d) Example of a U373 cell stably expressing AP2-eGFP (green) growing on an adhesive micropattern of Atto633-labelled PLL-g-PEG (red). Adhesive (black) and non-adhesive (red) areas separated by the PLL-g-PEG signal for further analysis of CME dynamics. Scale bars: 10 µm. (e) Lifetime versus maximal fluorescence intensity of AP2-eGFP of CME tracks from adhesive (black) and non-adhesive (black) areas of a representative cell. Each dot represents one tracked CME event of a 10 minute long movie. Persistent clathrin-coated plaques are marked by a grey circle. Number of tracks at the attached plasma membrane part: 2,849, at the non-attached plasma membrane part: 87.

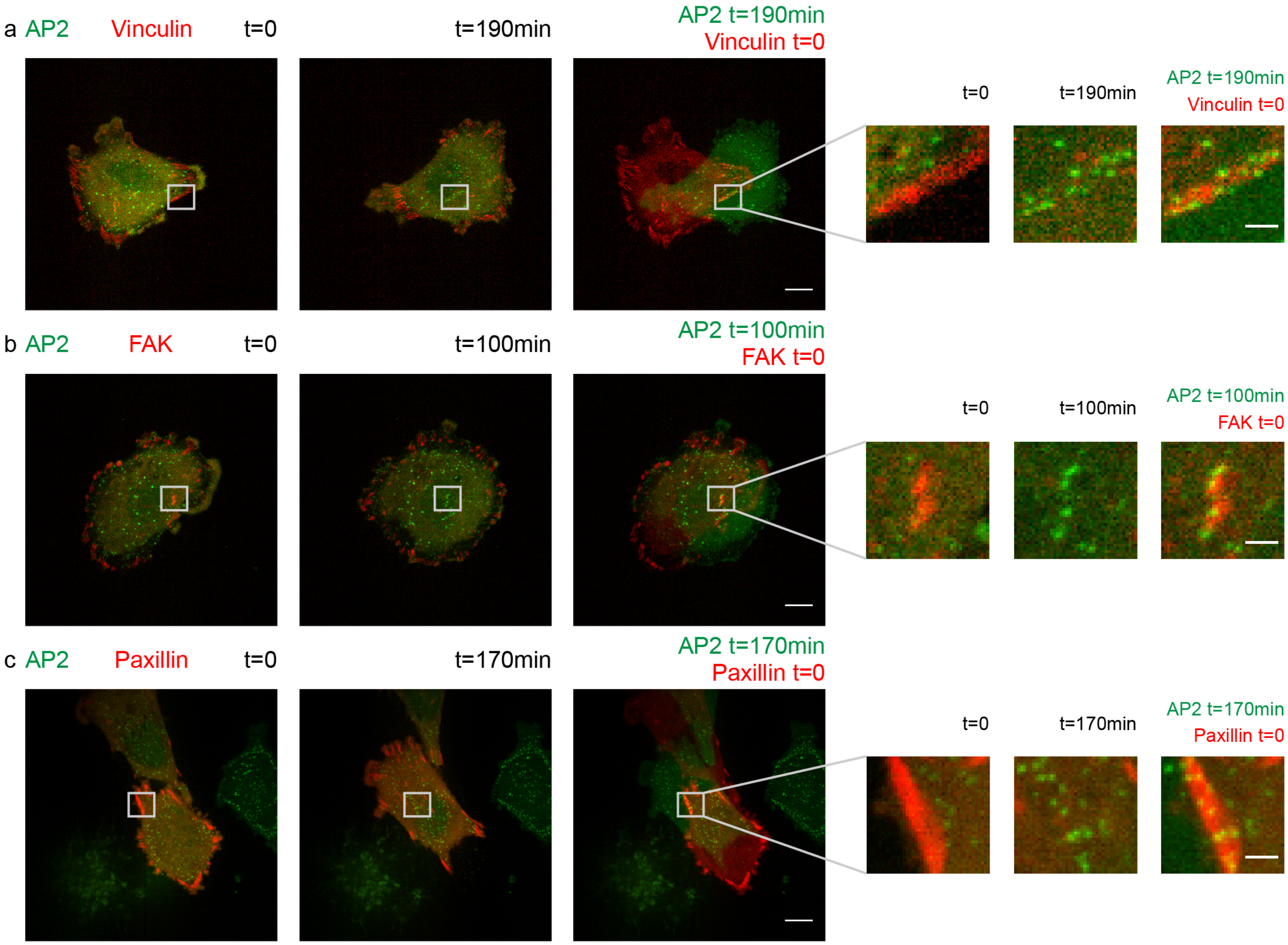


**Supplementary Figure 3: Different FA markers show switch from FAs to clathrin-coated plaques during cell migration.**

Live-cell confocal spinning disc microscopy of U373 stably expressing AP2-eGFP (green) and transiently expressing FA-associated proteins fused to the fluorescent protein mCherry (red): mCherry-vinculin (a), mCherry-FAK (b), or mCherry-paxillin (c). Left: Overview of a representative migrating cell at time 0 and a later time point (190, 100, or 170 minutes, respectively). Merged images of the FA protein signal at time 0 (red) and the AP2-eGFP signal at the later time point (green). Right: Zoom on FAs that switch to clathrin-coated plaques. Scale bars: 10 µm (overview), 2 µm (zoom).

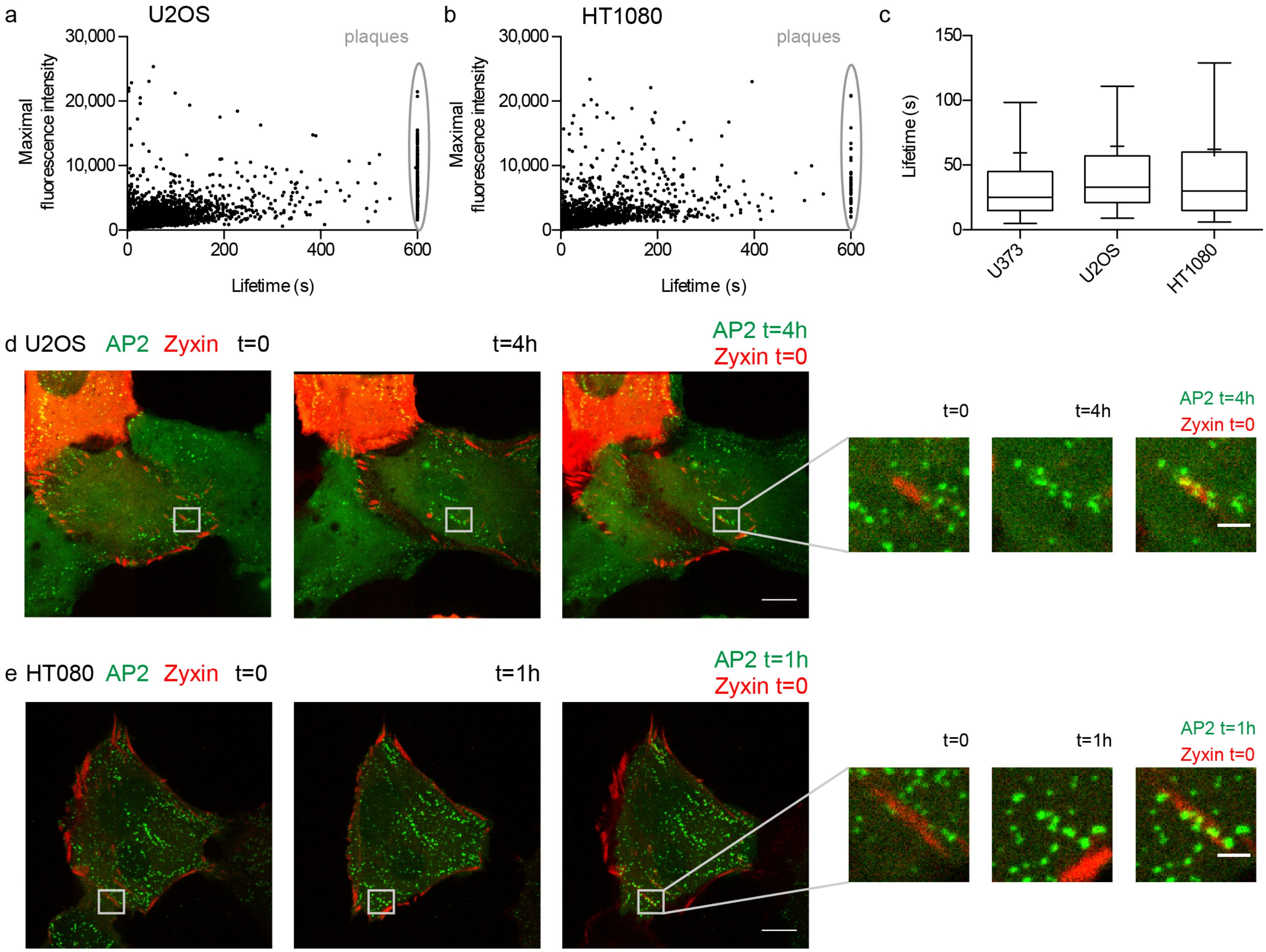


**Supplementary Figure 4: Switch from FAs to clathrin-coated plaques in U2OS and HT1080 cells.**

Lifetime versus maximal fluorescence intensity of AP2-eGFP of CME tracks for U2OS (a) and HT1080 (b). Shown are plots of a representative cell. Each dot represents one CME event in a 10 minute long movie. Number of tracks: 5,492 (U2OS) and 3,287 (HT1080). (c) Box and whiskers plot showing the lifetime distributions of CME in U373, U2OS, and HT1080 computed from three to four cells for each cell line. Whiskers represent 10-90 percentile, box represents second and third quartile, the line marks the median, and the cross marks the mean. Number of tracks: 4,472 (U373), 19,366 (U2OS), and 7,144 (HT1080). Live-cell confocal spinning disc microscopy of U2OS (d) and HT1080 (e) stably expressing AP2-eGFP (green) and transiently expressing mCherry-zyxin (red). Left: Overview of a representative migrating cell at time 0 and a later time point (4 hours and 1 hour for U2OS and HT1080, respectively). Merged image of the mCherry-zyxin signal at time 0 (red) and the AP2-eGFP signal at the later time point (green). Right: Zoom on FAs that switch to clathrin-coated plaques. Scale bars: 10 µm (overview), 2 µm (zoom).

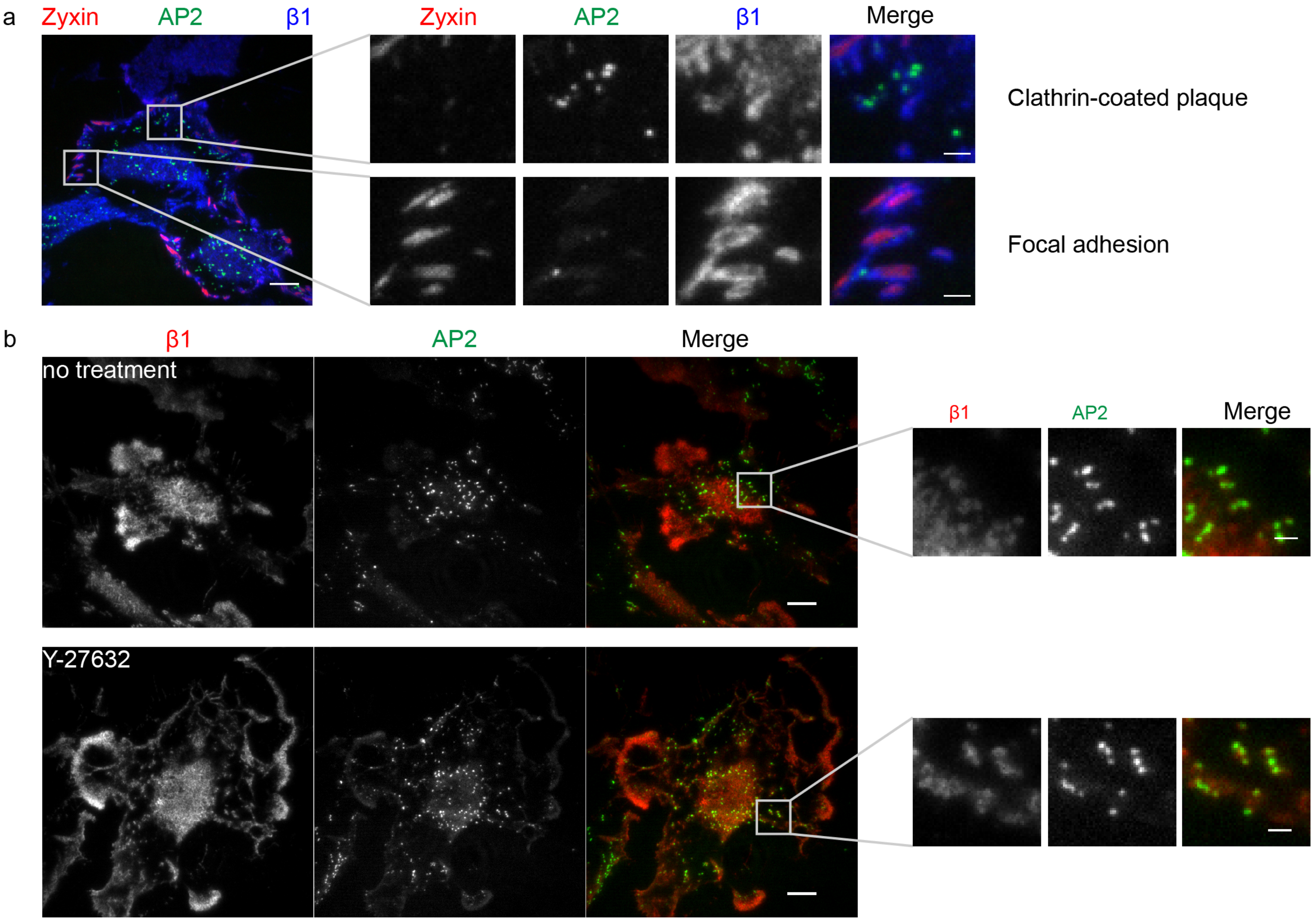


**Supplementary Figure 5: Enrichment of β1 integrin at clathrin-coated plaques.**

(a) Representative image from total internal reflection fluorescence (TIRF) microscopy of U373 stably expressing AP2-eGFP (green) and transiently expressing mCherry-zyxin (red) immunostained for β1 integrin (blue). Right: Zoom on clathrin-coated plaques marked by bright AP2-eGFP signal (upper panel) and on FAs marked by mCherry-zyxin (lower panel). (b) Representative image from TIRF microscopy of U373 stably expressing AP2-eGFP (green) immunostained for integrin β1 (red) untreated (upper panel) or treated with Y25632 (10 μM) for 30 minutes (lower panel). Right: Zoom on clathrin-coated plaques. Scale bars: 10 μm (overview), 2 μm (zoom).

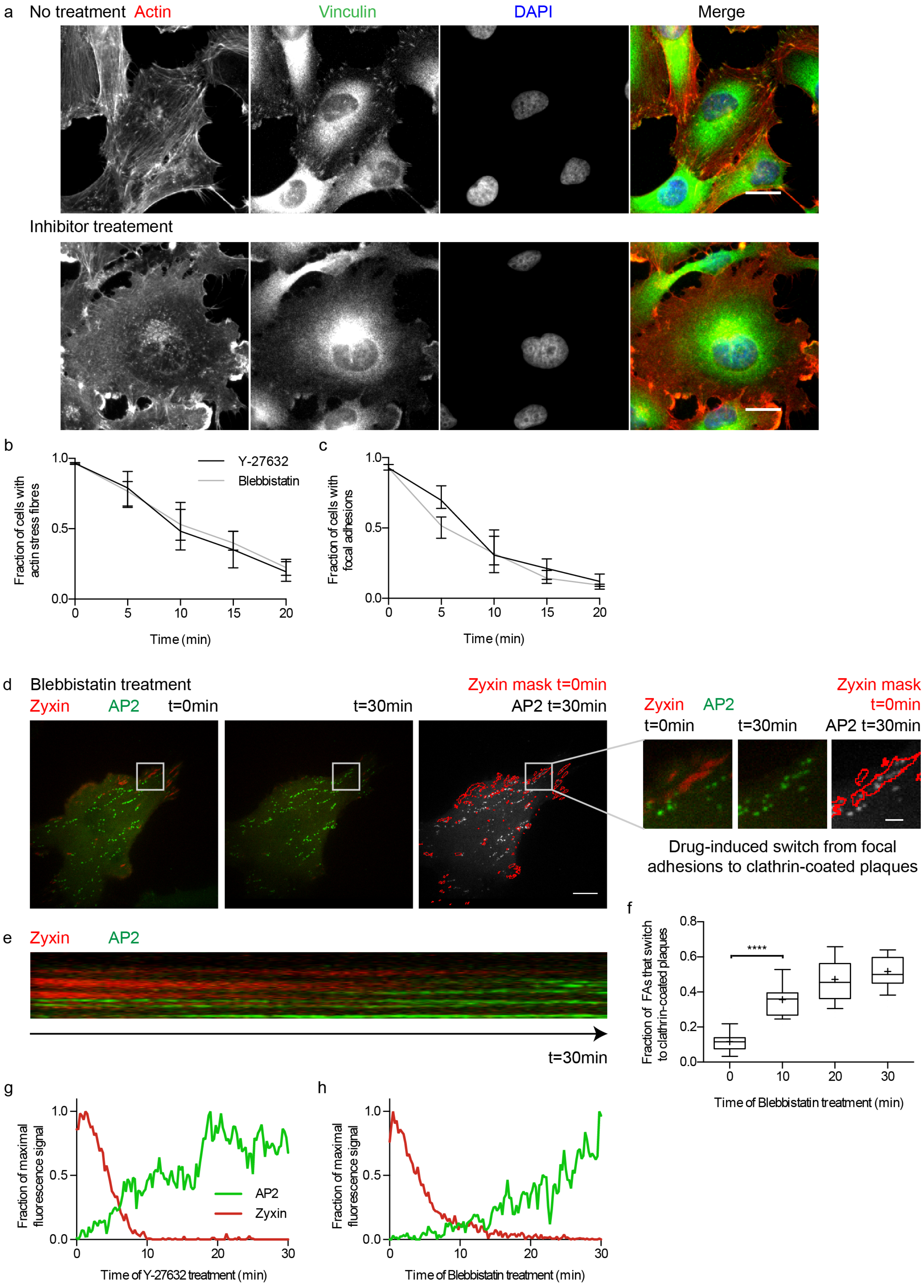
a
No treatment
Actin
Vinculin
DAPI
Merge
Inhibitor treatement
b
Fraction of cells with actin stress fibres
Y-27632
Blebbistatin
Time (min)
c
Fraction of cells with focal adhesions
Time (min)
d
Blebbistatin treatment
Zyxin
AP2
t=0min
t=30min
Zyxin mask t=0min
AP2 t=30min
Drug-induced switch from focal adhesions to clathrin-coated plaques
e
Zyxin
AP2
t=30min
f
Fraction of FAs that switch to clathrin-coated plaques
****
Time of Blebbistatin treatment (min)
g
Fraction of maximal fluorescence signal
AP2
Zyxin
Time of Y-27632 treatment (min)
h
Fraction of maximal fluorescence signal
Time of Blebbistatin treatment (min)

**Supplementary Figure 6: Drug-induced FA disassembly leads to switch to clathrin-coated plaques.**

(a) Representative images of widefield microscopy of untreated (upper panel) and inhibitor treated (lower panel) U373 cells stained for actin with Alexa Fluor 647-labelled phalloidin (red), vinculin (green) and DNA with DAPI (blue). Scale bar: 10 µm. Kinetics of stress fibre (b) and FA (c) disassembly by treatment with Y-27632 (10 µM, black) or Blebbistatin (20 µM, grey). The graph shows the mean with SD of each time point. The results are computed from three independent experiments with over a hundred cells per experiment. (d) Live-cell confocal spinning disc microscopy of representative U373 cell stably expressing AP2-eGFP (green) and transiently expressing mCherry-zyxin (red) treated with Blebbistatin (20 µM). Cell before (left) and 30 minutes after drug treatment (middle). (Right) Merged images of a mask marking the mCherry-zyxin objects before (red) and the AP2-eGFP signal after Blebbistatin treatment (green). Right: Zoom on FA that switches to clathrin-coated plaques during the treatment. (e) Kymograph of the switch from FAs to clathrin-coated plaques shown in d over 30 minutes of Blebbistatin treatment. (f) Quantification of the switch from FAs to clathrin-coated plaques during Blebbistatin treatment. Whiskers represent 10-90 percentile, box represents second and third quartile, line marks the median and cross the mean. Results are computed from three repetitions. Statistical analysis: t test, n=30, P<0.01. Normalized fluorescence intensity profiles of AP2-eGFP (green) and mCherry-zyxin (red) of a representative switch from FA to clathrin-coated plaques during Y-27632 (g) and Blebbistatin (h) treatment.

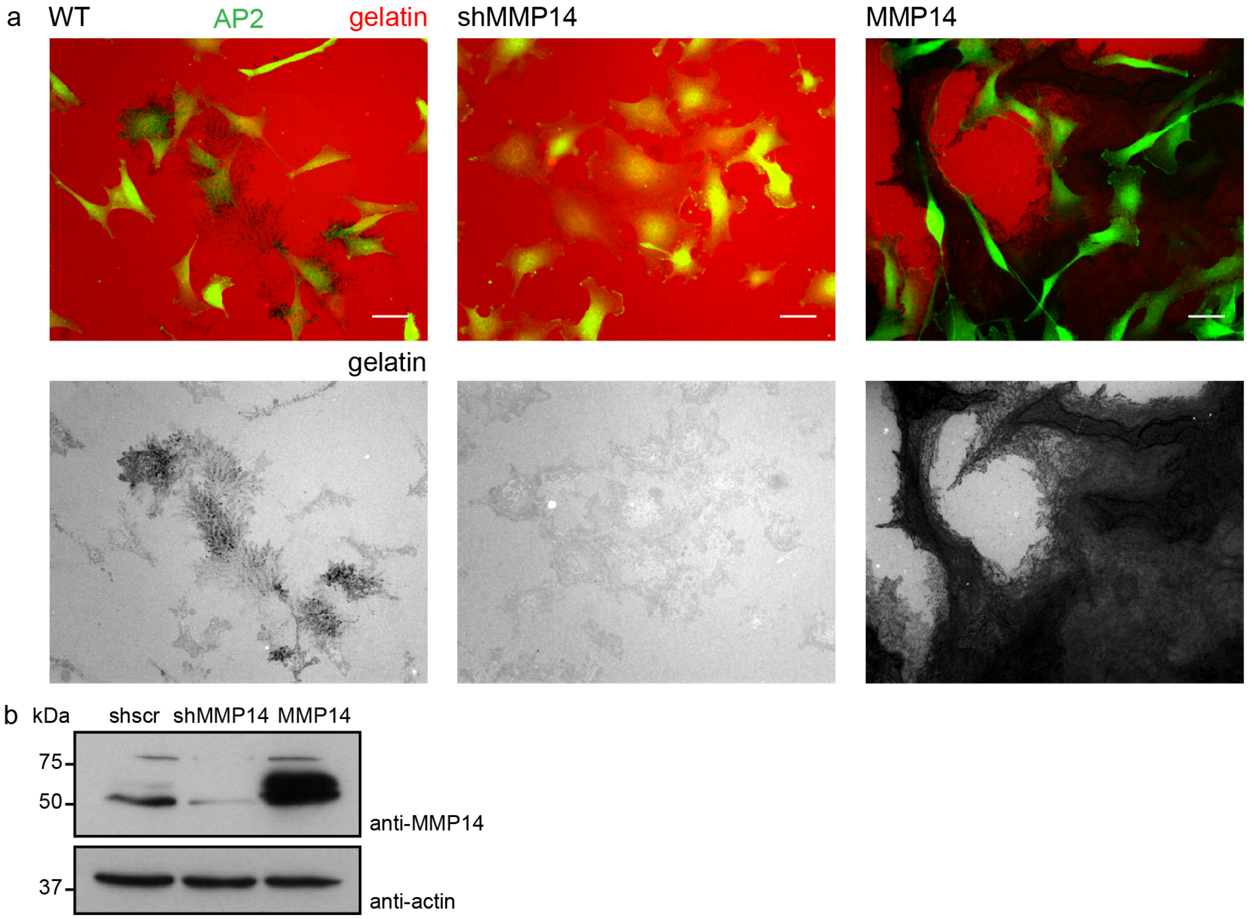


**Supplementary Figure 7: Gelatin digestion of U373 depends on MMP14.**

(a) Representative images of widefield microscopy of U373 stably expressing AP2-eGFP (green) wild-type (WT), stable knock down for MMP14 (shMMP14), or stable overexpression of MMP14 (MMP14) on coverslips coated with Alexa Fluor 647-labelled gelatin (red). Scale bar: 10 µm. (b) Western blot showing the protein level of MMP14 of U373 cell lines stably expressing shscrambled (shscr), shMMP14, and overexpressing MMP14. The protein levels of actin were used as loading control.

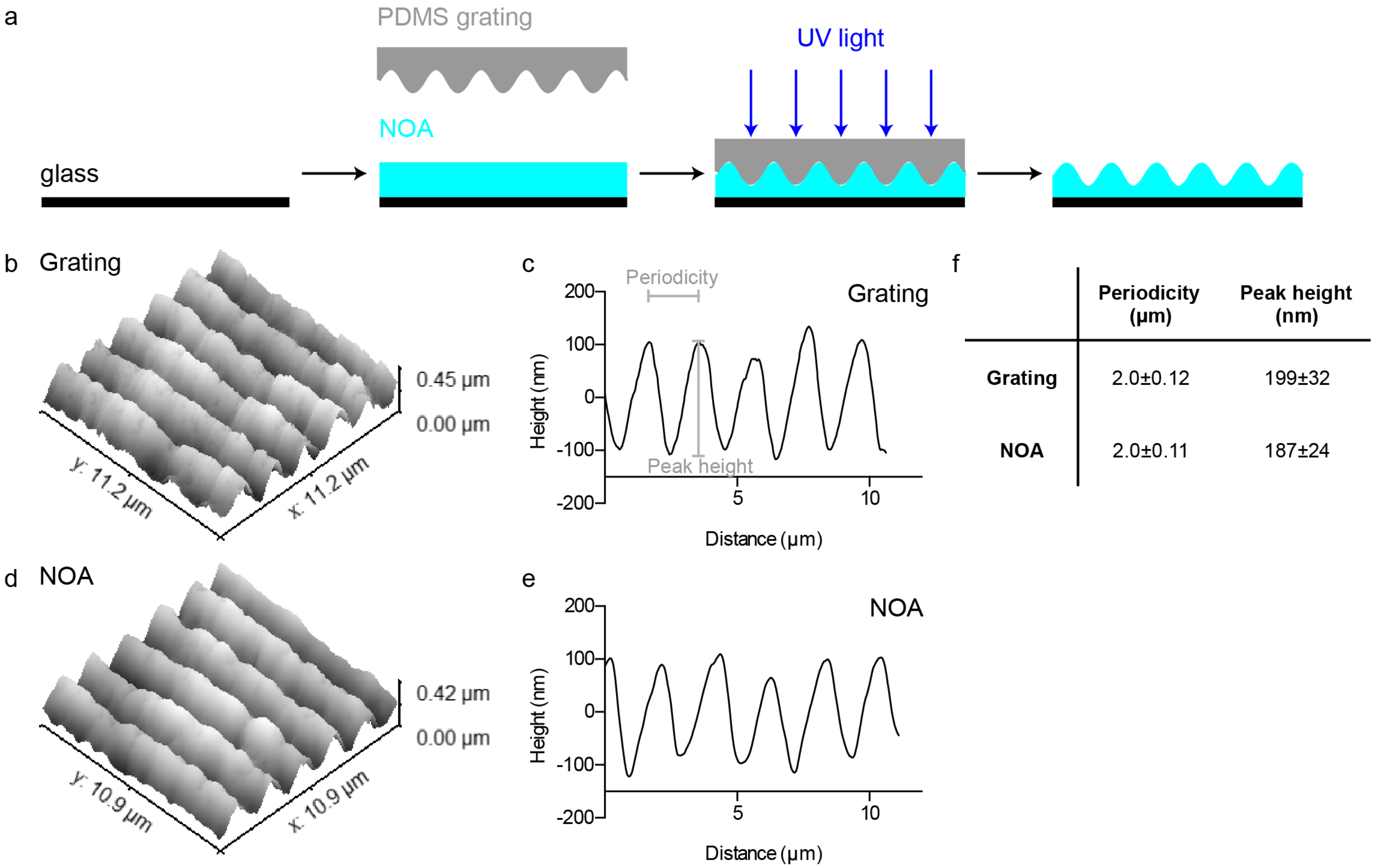


**Supplementary Figure 8: Production of optically clear 3D-micropattern.**

(a) Schematic illustrates the production of optically clear 3D-micropatterns. First, a drop of optically clear adhesive (NOA, turquois) was added on a glass coverslip and sandwiched by a PDMS grating (grey) generated from a diffraction grating. The optically clear adhesive was cured by UV-irradiation and finally, the PDMS grating was peeled from the optically clear 3D-micropatterns. Comparison of the topography of the diffraction gratings (b-c) and the optically clear 3D-micropatterns (d-e) by atomic force microscopy (AFM). 3D-representation of the topography of diffraction gratings (b) and optically clear 3D-micropatterns (d). (c and e) Line profile of the topography shown in b and d. Periodicity and peak height are marked. (f) Table lists the average periodicity and the peak height of diffraction gratings and optically clear 3D-micropatterns. Shown are the mean with SD, computed from three AFM measurements, with n=25 (periodicity) and n=28 (peak height).

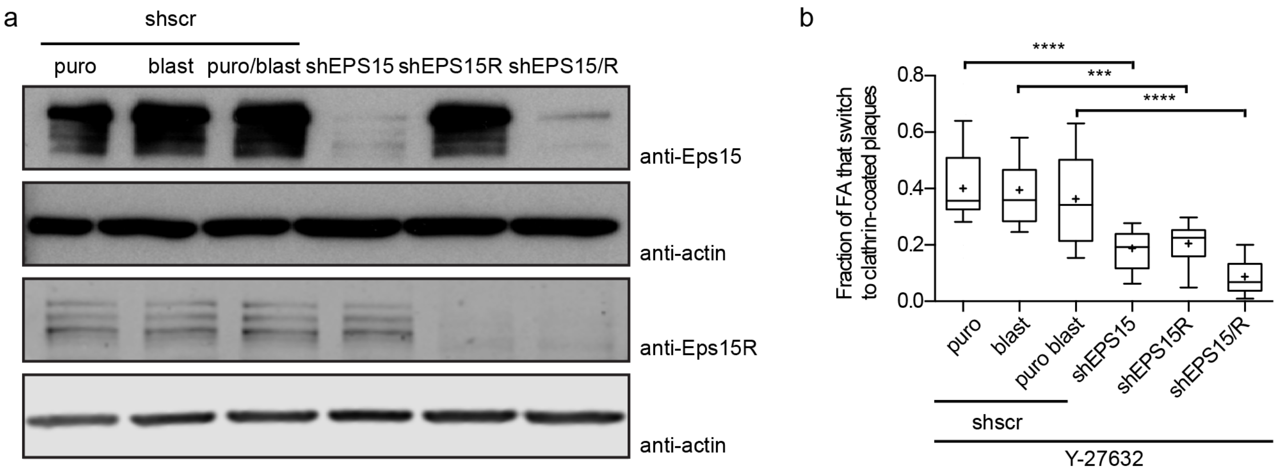


**Supplementary Figure 9: Eps15/R depletion reduces the efficiency of the switch from FAs to clathrin-coated plaques.**

(a) Western blot showing the protein level of Eps15 (top) and Eps15R (bottom) of U373 cell lines stably expressing shscrambled (shscr), shEPS15, shEPS15R and shEPS15 together with shEPS15R (shEPS15/R). The protein levels of actin were used as loading control. (b) Normalized quantification of the switch from FAs to clathrin-coated plaques after 20 minutes of Y-27632 treatment of U373 shEPS15, shEPS15R and shEPS15/R. Data was normalized to the mean of U373 stably expressing the corresponding shscr. Shown is the mean with SD computed from three repetitions. Statistical analysis: t test, n=28 (shscr puro), n=30 (shscr blast), n=24 (shscr puro blast), n=30 (shEPS15), n=25 (shEPS15R), n=29 (shEPS15/R), $P<0.01$.

### ***Supplementary References***


1. Bucher, D. *et al.* Clathrin-adaptor ratio and membrane tension regulate the flat-to-curved transition of the clathrin coat during endocytosis. *Nat. Commun.* **9,** 1109 (2018).

2. Pitaval, A., Christ, A., Curtet, A., Tseng, Q. & Théry, M. in *Cilia, Part B* (ed. Marshall, W. F. B. T.-M. in E.) **525,** 109–130 (Academic Press, 2013).
